# The effect of environmental variation on the diversity and composition of the three-spined stickleback microbiome

**DOI:** 10.64898/2026.05.06.723255

**Authors:** F Gyapong, M Barnes, B Fisher, T Guetta-Baranes, A MacColl, FJ Whelan

## Abstract

The fish skin microbiome serves as a protective barrier, influencing host health and facilitating interactions between the host and its environment. While several studies have characterised the composition and roles of the fish skin microbiome, there remains a paucity of data on how environmental variation influences these microbes in natural populations. Here, we used 16S rRNA gene sequencing to characterise the skin microbiome of wild three-spined stickleback populations and examine how environmental factors influence microbial diversity and community composition across 17 freshwater lochs on the island of North Uist, Scotland. Analysis of 239 samples revealed a set of dominant bacterial genera commonly associated with other fish species, including *Janthinobacterium*, *Pseudomonas*, *Acinetobacter*, and *Psychrobacter,* that constituted a core skin microbiota across lochs. Microbiome composition was primarily shaped by environmental variables, particularly habitat, water pH, conductivity, and metal concentrations, with pH emerging as a key driver of community structure. Host sex also influenced microbiome variation, with several taxa differing in relative abundance between males and females. Alpha-diversity was higher among stickleback fish from lochs with a neutral pH compared with those from alkaline and acidic environments. Differential abundance analyses identified 27 and 24 amplicon sequence variants (ASVs), respectfully, associated with variations in pH and host sex, including members of *Psychrobacter, Sphingobacterium*, *Carnobacterium*, *Chryseobacterium*, and *Arthrobacter*, highlighting the combined influence of environmental and host factors on microbiome composition in wild fish populations in freshwater environments.

## Background

Host-associated skin microbes play a key role in their host’s physiology, ecology, adaptation, and evolution (Fischbach & Segre, 2016; Henry et al., 2021). In fish, the skin functions as a protective barrier and is covered by a mucus layer that harbours microbes and remains in continuous contact with the surrounding environment (Dash et al., 2018; Dodd et al., 2020). This microbial community, collectively known as the skin microbiome, facilitates homeostasis of host immunity, produces antimicrobial compounds, and prevents the colonisation of potential pathogens, thereby maintaining the health and stability of the fish skin ecosystem (Hu et al., 2021; Peatman et al., 2015; Ross et al., 2019; Sultana et al., 2022).

Emerging evidence suggests that the composition of the skin microbiome is influenced by both host-specific traits (e.g. sex, body size, age) and environmental factors (e.g. water chemistry, location, diet) (D. Rosado et al., 2021; Ross et al., 2019; Sehnal et al., 2021a). For example, the skin microbiome composition may vary with host age (D. Rosado et al., 2021) and is known to be affected by abiotic factors such as seasonal variation and pollution (Bierlich et al., 2018; Georgala, 1958; Hooper et al., 2019; Wilson et al., 2008) and biotic factors such as host dietary composition (Chiarello et al., 2018; Horlick et al., 2020; Landeira-Dabarca et al., 2013). Hence, understanding the dynamics that shape these microbes is essential, especially in wild fish populations exposed to complex environmental stressors.

Numerous studies across a range of study systems have highlighted the influence of both host traits and environmental factors on fish skin microbiome composition and diversity (A. G. Bell et al., 2024; Berggren et al., 2022; Horlick et al., 2020; Ross et al., 2019; Sehnal et al., 2021a). Collectively, these studies demonstrate that variation in the fish skin microbiome is shaped by host-specific traits, including life stage, body size, and sex, as well as by environmental conditions such as water chemistry, seasonal changes, and habitat characteristics. However, the relative contributions of these drivers and their interactions in natural fish populations remain poorly understood. For example, heavy metals such as copper, cadmium, zinc, lead, and chromium are pervasive pollutants in freshwater systems and have been shown to influence non-host associated microbiomes in freshwater sediments (e.g., mineral and organic particles accumulated at the bottom of lakes and rivers) and in biofilms on submerged substrates (e.g., rocks and aquatic vegetation; Aceves-Suriano et al., 2023; Hu et al., 2025; Li et al., 2020; Wen et al., 2024). The influence of these factors on the host-associated skin microbiome of wild fish remains incompletely characterised. In particular, few studies have simultaneously examined multivariate environmental gradients (e.g., pH, conductivity, and metal levels) alongside host traits, thereby limiting understanding of their combined effects on skin microbiome composition.

To address these knowledge gaps, we investigated the skin microbiome of the three-spined stickleback, *Gasterosteus aculeatus* (hereafter referred to as ‘stickleback’), a well-established model organism in evolutionary biology, ecology, and environmental monitoring (Bell & Foster, 1994; Catteau et al., 2020; Katsiadaki et al., 2006; MacColl et al., 2013; Magalhaes et al., 2016). In many populations worldwide, stickleback can be broadly divided into two main morphotypes: freshwater (smaller, slender bodies with reduced/complete loss of armour plates and pelvic spine) and marine/anadromous (larger, broader bodies with full armour plates and a pelvic spine; Bell & Foster, 1994; Shapiro et al., 2004; Wiig et al., 2016). The anadromous form migrates to coastal lagoons and freshwater habitats to reproduce, and the offspring return to the ocean, whereas the freshwater form remains in freshwater habitats year-round, inhabiting lakes, streams, rivers, and ponds (Aguirre et al., 2022; MacColl et al., 2013).

Stickleback have recently emerged as a model organism for microbiome studies owing to their broad geographic distribution, natural genetic variation, and ecological diversity (Milligan-Myhre et al., 2016; Shankregowda et al., 2023; Small et al., 2023). To date, microbiome research in sticklebacks has focused primarily on the gut, demonstrating associations with sex, habitat, and host genotype (Härer et al., 2025; Milligan-Myhre et al., 2016; Small et al., 2023). A key gap in stickleback microbiome studies is the absence of direct assessment of the relative contributions of environmental factors and host-specific traits to the composition and diversity of the skin microbiome.

Here, we address this gap by using freshwater-resident stickleback populations from the island of North Uist, Scotland, where lochs (lakes) exhibit geographical and ecological variation, ranging from acidic to neutral and alkaline conditions, as well as differences in conductivity and metal concentrations (MacColl et al., 2013). This setting provides a natural system for examining environmental influences on the skin microbiome while avoiding potential confounding effects of salinity and the life-history transitions associated with anadromous forms. Establishing this freshwater baseline will also facilitate future comparisons between freshwater resident and anadromous populations. North Uist has long been a site of scientific interest owing to its diverse lochs and genetically structured stickleback populations (Campbell, 1985; Giles, 1983; MacColl et al., 2013; Magalhaes et al., 2016; Waterston et al., 1979). Previous work has shown that stickleback populations across the island differ in traits such as resistance to parasitic infection, highlighting the ecological heterogeneity that makes this system well-suited for microbiome research (de Roij et al., 2011; MacColl & Chapman, 2010; Robertson et al., 2017).

Using 16S rRNA gene sequencing of the skin microbiome of wild stickleback populations from North Uist, we aimed to address the following research questions: (i) What is the diversity and composition of the stickleback skin microbiome on North Uist? (ii) How do environmental factors such as pH, conductivity, heavy metal concentrations, and population (i.e., loch identity), shape microbiome composition? and (iii) To what extent do host-associated variables, such as fish length and sex, influence microbiome diversity and composition? Together, these questions aim to disentangle the relative roles of environmental and host-associated factors in structuring the stickleback skin microbiome.

## Materials and Methods

### Study area

307 stickleback fish were sampled from 17 lochs (lakes) on the island of North Uist, Scotland over 5 days during a single field season (57°35′N; 7°18′W; **Figure 1**). These lochs are isolated from one another in small catchment areas, vary substantially in size, and are generally shallow, with an average depth of 2.8 m (Giles, 1983; MacColl et al., 2013; Waterston et al., 1979). The island has an unusual surface geology that results in differing water chemistry and is associated with substantial variation in stickleback body size and morphology (Giles, 1983; MacColl et al., 2013). The east side of the island is characterised by acidic, oligotrophic lochs, whereas the west side hosts mesotrophic to eutrophic lochs (Figure 1) (MacColl et al., 2013; Waterston et al., 1979). Based on measured pH values across the island, lochs were grouped into three categories: acidic (pH < 7.0), neutral (pH 7.0–7.40), and alkaline (pH > 7.40; **Figure 1**, **Table 1**).

**Figure 1:**
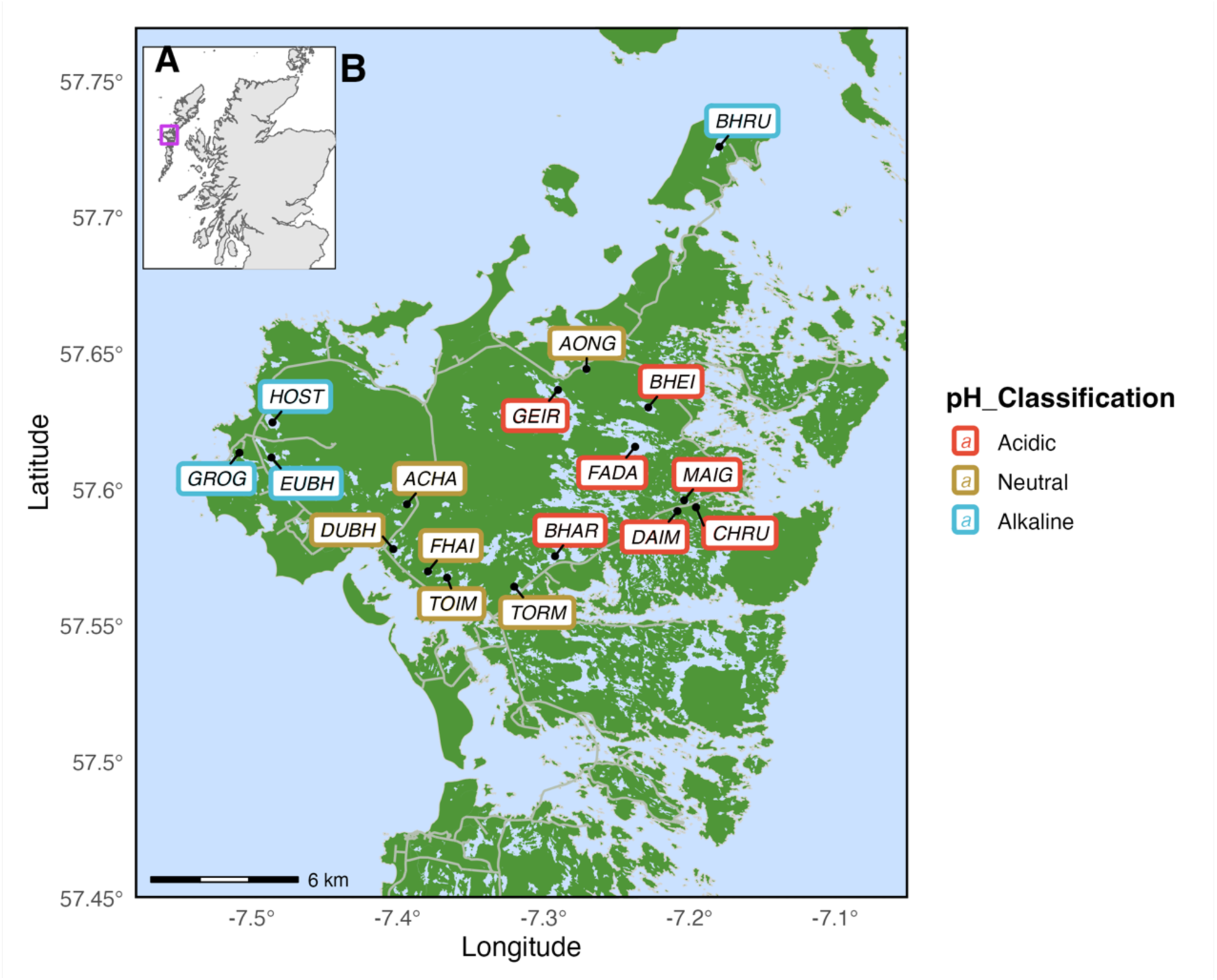
Geographical location and pH classification of study sites in North Uist. A: Map of Scotland indicating the study region of North Uist (highlighted with a purple boundary box). **B:** Detailed map of North Uist depicting the locations of sampled lochs. Site labels are colour-coded by pH classification: Acidic (**red**), Neutral (**gold**), and Alkaline (**blue**).

**Table 1:** Summary of the freshwater sticklebacks and metadata variables.

| Loch (code) | Samples collected | Samples sequenced | Mean ± SD length (mm) | Median length (mm) | Sampling date | pH | pH classification |
| --- | --- | --- | --- | --- | --- | --- | --- |
| a'Bharpa (**Bhar**) | 18 | 17 | 33.4 ± 3.41 | 33 | 11/05/2024 | 6.30 | Acidic |
| Fada (**Fada**) | 11 | 10 | 33.2 ± 3.76 | 33.2 | 11/05/2024 | 6.58 | |
| an Daimh (**Daim**) | 20 | 12 | 34.9 ± 3.87 | 33.5 | 11/05/2024 | 6.6 | |
| Bheireagvat (**Bhei**) | 20 | 18 | 30.3 ± 3.98 | 30.5 | 11/05/2024 | 6.74 |  |
| nan Geireann (**Geir**) | 7 | 5 | 29.8 ± 4.66 | 27 | 12/05/2024 | 6.81 |  |
| Chadha Ruaidh (**Chru**) | 20 | 18 | 42.1 ± 3.21 | 43 | 11/05/2024 | 6.83 |  |
| Maighdein (**Maig**) | 17 | 14 | 30 ± 4.44 | 31 | 11/05/2024 | 6.97 |  |
| Dubhasaraidh (**Dubh**) | 20 | 16 | 38.6 ± 2.75 | 39 | 12/05/2024 | 7.05 | Neutral |
| Fhaing Buidhe (**Fhai**) | 20 | 19 | 32.1 ± 5.27 | 31 | 13/05/2024 | 7.05 |  |
| Aonghais (**Aong**) | 20 | 15 | 29.5 ± 2.45 | 29 | 12/05/2024 | 7.1 |  |
| an Toim (**Toim**) | 20 | 19 | 33.4 ± 4.2 | 33 | 13/05/2024 | 7.16 |  |
| Tormasad (**Torm**) | 20 | 17 | 33.4 ± 4.15 | 33 | 13/05/2024 | 7.27 |  |
| a'Charra (**Acha**) | 20 | 17 | 37.8 ± 6.89 | 36.5 | 13/05/2024 | 7.34 |  |
| Eubhal (**Eubh**) | 20 | 16 | 38.6 ± 6.64 | 37 | 12/05/2024 | 7.86 | Alkaline |
| Grogary (**Grog**) | 10 | 6 | 34.8 ± 5.34 | 32.5 | 12/05/2024 | 8.10 |  |
| Hosta (**Host**) | 20 | 10 | 42.2 ± 3.71 | 41.5 | 12/05/2024 | 8.19 |  |
| Bhrusda (**Bhru**) | 10 | 10 | 37.4 ± 2.93 | 37.8 | 09/05/2024 | 8.44 |  |
| *Mean, median and SD± was calculated using the sequenced samples. |  |  |  |  |  |  |  |

### Sample collection

Stickleback fish were captured in the lochs of North Uist using Gee minnow traps (Dynamic Aqua, Vancouver, Canada) from 09-13 May 2024. Six traps were set for two consecutive nights at each loch. Upon retrieval, all sticklebacks captured from each loch were temporarily transferred to 10 L buckets containing water from the same loch prior to processing. At each site, up to 20 fish were opportunistically sampled for skin swabbing; however, at Bhru, Fada, Geir, Grog, and Maig, fewer than 20 fish were captured, and all individuals caught were swabbed (**Table 1**). The captured sticklebacks were handled using sterile nitrile gloves (StarGuard® Comfort, Nitrile Gloves, SG-C-M, Starlab UK) that had been disinfected with 70% ethanol; this procedure was repeated between fish to minimise cross-contamination. Each fish was rinsed with sterile distilled water to remove transiently associated environmental microbes. Each fish was gently restrained on top of a sterile foam surface moistened with sterile distilled water. A sterile cotton swab (ISO 18385, cotton, 150 mm, SARSTEDT LTD, UK) was used to collect the skin mucus by swabbing 10 times on the left and 10 times on the right lateral sides of the fish. Upon collection of the skin mucus, the swabs were immediately returned to labelled 1.5 mL Eppendorf tubes containing 400 µL of Copan Liquid Amies (Copan Liquid Amies Elution Swab, Italy). The tubes containing the swabs were stored at 4°C during fieldwork (within 48 h) before being transported to the laboratory, where they were stored at - 20°C for further processing. The swabbed fish were euthanised using an overdose of MS-222 (400 mg/L), followed by pithing, and their standard length was measured in mm. The sex of each fish was determined by visually inspecting the gonads through abdominal dissection; individuals with testes were identified as males, and those with eggs were identified as females.

### Water chemistry

Water parameters, including temperature, conductivity, and pH were measured from all lochs using a calibrated multi-parameter probe (Multi 340i, WTW, Weinheim, Germany). For metal analysis, 1 L of water from each loch was collected into sterile bottles. A 20 mL subsample was then drawn into a 20 mL sterile syringe (Terumo syringe, TERUMDSS20ESE, VWR International) and passed through a 0.2 μm sterile syringe filter (Sartorius Minisart NML Syringe Filters, 10527401; Fisher Scientific Ltd) into a 25 mL sterile container pre-filled with 5 mL of 10% nitric acid (HNO₃), resulting in a final acidified concentration of 2% HNO₃.

Samples were analysed using inductively coupled plasma mass spectrometry (ICPMS; Thermo Fisher Scientific iCAP-Q, Bremen, Germany), alongside known standards and operational blanks for comparison and quality control. Analyses of the elemental composition were carried out at the ICP-MS facility located in the Gateway Building, Sutton Bonington Campus, University of Nottingham, UK.

### DNA extraction

Genomic DNA was extracted from 239 swab samples (**Table 1**), following previously described protocols (Whelan et al., 2020). Briefly, in a 2 mL plastic screw-top tube containing 0.2 g of 0.1 mm glass beads (Mo Bio Laboratories, 13118-50), 800 µL of 200 mM monobasic NaPO4 (pH=8) and 100 μL of guanidine thiocyanate-EDTA-Sarkosyl were added, followed by the addition of 300 µL of the Copan amies solution that the swab has been collected into. Cells were lysed using a bead beater (Precellys Evolution; EQ02520-300-RD000.0; Bertin Technologies) at 4500 rpm for 1 min. Subsequently, 50 µL of lysozyme (100 mg/mL) (Sigma-Aldrich, L6876-10G) and 10 µL of RNase A (10 mg/mL in H₂O; Qiagen, 19101) were added, and the mixture was vortexed and incubated at 37°C for 1.5 h.

Following, 25 µL of 25% SDS (diluted in ddH₂O), 25 µL of Proteinase K (Sigma-Aldrich, P2308-1G), and 62.5 µL of 5 M NaCl (filter sterilised) were added to the samples, vortexed, and incubated at 65°C for 1 h. The screw-cap tubes were centrifuged at 13,500g for 5 min. After centrifugation, 900 µL of the supernatant was transferred to 2 mL Eppendorf tubes containing 900 µL of a 25:24:1 phenol-chloroform-isoamyl alcohol solution (Sigma-Aldrich, P3803-400mL), yielding an equal-volume sample-to-phenol-chloroform mixture.

The mixture was vortexed and centrifuged at 13,000g for 10 min. After centrifugation, the top layer was carefully transferred to a new 1.75 mL Eppendorf tube pre-filled with 200 µL of DNA-binding buffer (DNA Clean and Concentrator-25, D4034, Zymo Research) and mixed. This solution was transferred to a DNA column (DNA Clean and Concentrator-25, D4034, Zymo Research) in 600 µL aliquots. The columns were spun at 12,000g for 1 min, and the flow-through was discarded.

After the sample had passed through the column, 200 µL of wash buffer (DNA Clean and Concentrator-25, D4034, Zymo Research) was added to the column, which was then spun at 12,000g for 1 min, and the flow-through was discarded; this process was repeated twice. The columns were transferred to new, sterile 1.75 mL Eppendorf tubes, and 50 µL of sterile DNase/RNase-free ddH₂O preheated to 65°C was added to the centre of each column. The columns were incubated at room temperature for 5 min, and the DNA was eluted into Eppendorf tubes by centrifugation at 12,000g for 1 min. Extracted DNA was quantified using a Qubit 4 Fluorometer (Invitrogen, Thermo Fisher Scientific, Q33226) and stored at -20°C. Two negative controls were included at the extraction stage: (i) an aliquot of Copan Liquid Amies transport medium to assess potential contamination from the collection medium, and (ii) an extraction blank containing all reagents to monitor potential contamination introduced during the extraction workflow. These controls were processed concurrently with the samples throughout all subsequent steps.

### Polymerase Chain Reaction and 16S rRNA gene sequencing

The universal primers to the 16S rRNA gene variable 3 and 4 regions derived from those described in (Apprill et al., 2015; Bartram Andrea et al., 2011; Herlemann et al., 2011) were used to amplify the variable 3 and 4 regions of the 16S rRNA gene. The 341F and 806R 16S rRNA gene primers, with unique six-base-pair barcodes incorporated into both the forward and reverse primers to enable multiplex amplification, yielding a 601-bp amplicon (Apprill et al., 2015; Herlemann et al., 2011). A 50 µL PCR reaction was prepared for each sample (as previously described, Whelan et al., 2020), split into three equal volumes for amplification. After PCR, the three reactions were combined into a single pooled sample (50 µL) prior to sequencing. The PCR reaction comprised: 5 µL of reverse and forward primers (1 µM), 1.5 µL MgCl₂ (50 mM), 1 µL dNTPs (10 mM), 0.25 µL Taq polymerase (5 U), 5 µL 10× buffer, 2 µL BSA (10 mg/mL), 30 ng DNA template, and nuclease-free water up to a final volume of 50 µL. The amplification conditions consisted of an initial denaturation step at 94°C for 5 min, followed by 5 cycles of 94°C for 30 s, 47°C for 30 s, and 72°C for 40 s. This was followed by 25 additional cycles of 94°C for 30 s, 50°C for 30 s, and 72°C for 40 s, with a final step at 72°C for 10 min. The presence of a PCR product was confirmed by electrophoresis on a 1.5% agarose gel; only samples with visible bands were submitted for sequencing.

PCR products, including negative controls, were sequenced on the AVITI short-read platform (2 × 300 bp paired-end) at DeepSeq, University of Nottingham. Sequenced controls included: (i) Copan Liquid Amies transport medium processed through DNA extraction and PCR, (ii) DNA extraction blanks containing all reagents but no biological material, and (iii) PCR blanks consisting of all PCR reagents with no DNA. Libraries were purified using AMPure bead clean-up on an Opentrons robotic platform prior to quality control. The concentrations of the purified libraries were determined using a Qubit 4 Fluorometer with the DNA High Sensitivity Assay (Thermo Fisher Scientific), and the library profiles were assessed using the Agilent 4200 Tape Station with the High Sensitivity D5000 DNA Assay (Agilent Technologies). Libraries were pooled in equimolar amounts (one per 96-well plate) and quantified using the KAPA Library Quantification Kit for Illumina (Roche; KK4824). The pools were then merged into a final library prior to sequencing. The final pool was sequenced on the Element Biosciences AVITI platform using a 2 × 300 bp Cloudbreak FS Medium Output kit to generate paired-end reads.

### Processing of 16S rRNA gene sequencing data

Raw 16S rRNA gene sequencing data were generated across two independent AVITI sequencing runs. Sequencing data were pre-processed using Cutadapt (v5.1; Martin, 2011) to remove adapters and PCR primers, and to filter reads with a minimum Phred quality score of 20 and a minimum length of 100 bp. Amplicon sequence variants (ASVs) were inferred using the DADA2 pipeline (Callahan et al., 2016). To minimise potential batch effects associated with sequencing runs, error learning and denoising were performed separately for each run prior to merging ASV tables in DADA2 for downstream analyses. Chimeric sequences were identified and removed using the consensus method implemented in DADA2. Taxonomic classification of ASVs was performed using the SILVA reference database (v 138.2; Callahan, 2024; Quast et al., 2013). The six negative controls (DNA extraction blanks, Copan liquid Amies transport medium, and PCR controls) processed alongside the biological samples yielded very low bacterial reads: five controls had <10 reads, and the remaining control yielded 565 reads. Given the relatively low microbial biomass associated with fish skin samples (Fierer et al., 2025), potential contaminants were investigated using the prevalence-based method implemented in the R package *decontam* (v1.22.0), and no ASVs were classified as contaminants. Because samples were processed across two sequencing runs, potential batch effects were evaluated using PERMANOVA on Bray–Curtis and Aitchison distance matrices; microbiome composition did not differ significantly between sequencing runs (*p* > 0.05) (**Supplementary Figure 1**).

### Data analysis of 16S rRNA gene sequencing data and metadata

Following the described quality control measures, a total of 239 samples were included in the microbiome analysis. To identify any potential correlations among metadata variables and avoid reporting indirect associations with the microbiome composition, pairwise tests were performed using chi-squared tests (categorical variables), ANOVA (continuous vs categorical), or Spearman correlation (continuous vs continuous), as appropriate. When statistical tests resulted in p-values < 0.05, the null hypothesis that the variables were independent was rejected. A full list of tested variable pairs and corresponding statistics is provided (**Supplementary Table 1**).

All statistical analyses were conducted in R v4.3.2 (R Core Team, 2023), using phyloseq v1.28.0 (McMurdie & Holmes, 2013) and vegan v2.6.10 (Oksanen, 2022) packages. Alpha diversity was calculated using the Shannon (Shannon, 1948) and Simpson (Simpson, 1949) indices. Beta diversity was assessed using both the Bray–Curtis distance (Bray & Curtis, 1957) in phyloseq and the Aitchison distance (Aitchison, 1982) via the microbiome package v1.24.0 (Lahti & Shetty, 2019). For Bray–Curtis distances, data were rarefied to the minimum sequencing reads per sample in the dataset (n=12,123; **Supplementary file S1: Figure 1**). Aitchison distances were calculated on unrarefied counts using a centred log-ratio (CLR) transformation to maintain compositional integrity of the data.

Linear mixed models were used to test for associations between metadata variables (as fixed effects) and Shannon diversity, with loch (habitat) included as a random effect. Models were fitted using the lmer function from the lme4 package v1.1.36 (Bates et al., 2015), with p-values derived using the lmerTest package v3.1.3 (Kuznetsova et al., 2017). A Type III ANOVA was then used to assess the statistical significance of model terms, using the anova function from the car package v3.1-3 (Fox & Weisberg, 2019). The homogeneity of variance was assessed using Levene’s test (Fox & Weisberg, 2019), and if the test indicated unequal variances across lochs (p < 0.05), a non-parametric Kruskal–Wallis test (Kruskal & Wallis, 1952) was used to determine whether alpha diversity differed between lochs.

To investigate the contributions of metadata variables to variation in skin microbiome composition among sticklebacks, a permutational multivariate analysis of variance (PERMANOVA) test was employed, utilising the adonis2 function from the vegan package v2.6.10 (Oksanen, 2022). A full additive model was used, including the following explanatory variables: sampling day, sex, length, pH classification, conductivity, zinc (Zn), copper (Cu), lead (Pb), and loch. Both Bray–Curtis and Aitchison distances were used to evaluate beta diversity patterns and assess the significance of each term. Differential abundance of ASVs was determined across pH classifications (alkaline, acidic, and neutral) and sex (male and female) using the DESeq2 package v1.42.1(Love et al., 2014). For both analyses, p-values were adjusted for multiple testing using the Benjamini–Hochberg (BH) (Benjamini & Hochberg, 1995) method implemented in DESeq2, with a significance threshold set at an adjusted p-value < 0.05.

## Results

### The composition of the stickleback skin microbiome across lochs

The stickleback skin microbiome composition varies across lochs (**Figure 1a**). Across the 239 samples analysed, we identified 285 unique genera, with 182 having a cumulative relative abundance of ≥ 0.01% and 38 having a relative abundance of ≥ 1% in at least one sample. Each sample contains, on average, 22 genera; however, this ranged from 8 to 72 (SD ± 10). Lochs Toim and Torm exhibited the highest number of mean genera and greatest sample-level variation (32 ± 15 and 30 ± 18, respectively), whereas Bhei had the lowest number of mean genera (17 ± 5) (**Supplementary Table 2**). On average, samples from neutral pH lochs exhibited the highest mean number of genera (26 ± 12.2), followed by alkaline (24 ± 6.8), and acidic (18 ± 5.5).

Across all samples*, Janthinobacterium* was the most abundant genus, accounting for over 50% of the bacterial community, with a mean relative abundance of 49.87% across all samples (median 52.34%; range = 3.51–91.02%; **Supplementary Table 3**). Other abundant genera included *Pseudomonas* (mean relative abundance 22.70%, median 20.11%, range 4.19–62.73%), *Acinetobacter* (mean relative abundance 7.36%, median 3%, range 0.08–53.33%), and *Psychrobacter* (mean relative abundance 6.74%, median 2.13%, range 0.01–76.17%). These four genera were present in all samples. Additionally, *Exiguobacterium*, *Chryseobacterium, Carnobacterium, Comamonas,* and *Rahnella* were present at varying relative abundance, while a diverse group of taxa occurred at <1% relative abundance (**Figure 1a**). Variations in genus-level composition were found across different lochs. For example, *Arthrobacter* was dominant among sticklebacks in Toim and Torm, while Bhar and Bhru had a higher relative abundance of *Rahnella*.

To investigate the contributions of specific amplicon sequence variants (ASVs) within the microbiome, we analysed the ASV-level diversity within the four most abundant bacterial genera across individual stickleback samples. Here, we defined an ASV as a unique DNA sequence recovered from 16S rRNA gene amplicon sequencing that represents a bacterial variant at the species or strain level (see Methods; Fasolo et al., 2024). *Pseudomonas* exhibited the highest within-genus ASV diversity, with notable variation at the individual and loch levels (**Figure 1b**). ASV2 and ASV5 contributed the highest proportion of relative abundance of *Pseudomonas* in most samples, while several additional ASVs were present at lower relative abundance. Sticklebacks from Bhar and Bhru exhibited diverse *Pseudomonas* ASVs yet had relatively low genus-level abundance (<20%), whereas those from Chru, Dubh, and Eubh had higher genus-level abundance (> 40%). We observed a shift in the dominant ASVs which corresponded to pH classification (acidic, neutral, alkaline); for instance, ASV22 was more prominent in acidic lochs compared with neutral and alkaline lochs.

The ASV diversity within the *Acinetobacter* genus was similar to that of *Pseudomonas*, with variation across sites (**Figure 1c**). ASV9 was the major variant across many samples, while other ASVs (e.g., ASV10, ASV12, ASV14, ASV17, and ASV18) were present at lower, variable proportions. Overall, the relative abundance of *Acinetobacter* and its ASV diversity were low in most acidic and alkaline lochs, except Bhar and Bhru. In these lochs, most individuals exhibited a diversity of ASVs and had <20% relative abundance of the genus overall. In contrast, several samples from neutral lochs, particularly Fhai, Toim, Torm, and Acha, revealed greater ASV diversity, with some individuals exceeding 40% relative abundance of the genus.

*Psychrobacter* was dominated by a few ASVs that varied substantially among individuals and lochs (**Figure 1d**). ASV8, ASV11, and ASV15 were the predominant variants across most individuals. Psychrobacter ASVs were detected in sticklebacks from acidic, neutral, and alkaline lochs; however, this genera’s proportional contribution varied among these groups. In contrast to the pattern observed for *Acinetobacter* (**Figure 1c**), individuals from Bhar (acidic) and Bhru (alkaline) exhibited lower *Psychrobacter* abundance (<1%) and ASV diversity. Notably, these same lochs were the only acidic and alkaline lochs showing comparably higher *Acinetobacter* abundance and ASV diversity (**Figure 1c, 1d**).

The *Janthinobacterium* had the lowest ASV diversity among the four abundant genera (**Figure 1e**). It was dominated by a few highly abundant ASVs, with ASV1 contributing to the largest fraction in most samples. Samples from acidic lochs were characterised by a higher proportional contribution of ASV1, although ASV3 and ASV4 also contributed substantial fractions in several individuals. In contrast, samples from neutral lochs exhibited greater heterogeneity in ASV composition, with several ASVs contributing moderate proportions to the total *Janthinobacterium* abundance. Alkaline lochs showed a similar pattern of ASV1 dominance, but with increased contributions from additional ASVs and pronounced within-loch variability (e.g., individuals from Eubh and Host). Low-abundance ASVs (<1%) collectively contributed a small fraction of the total *Janthinobacterium* abundance across samples, and this contribution was lower than that observed for *Pseudomonas* and *Acinetobacter* (**Figure 1b, c**).

**Figure 1:**
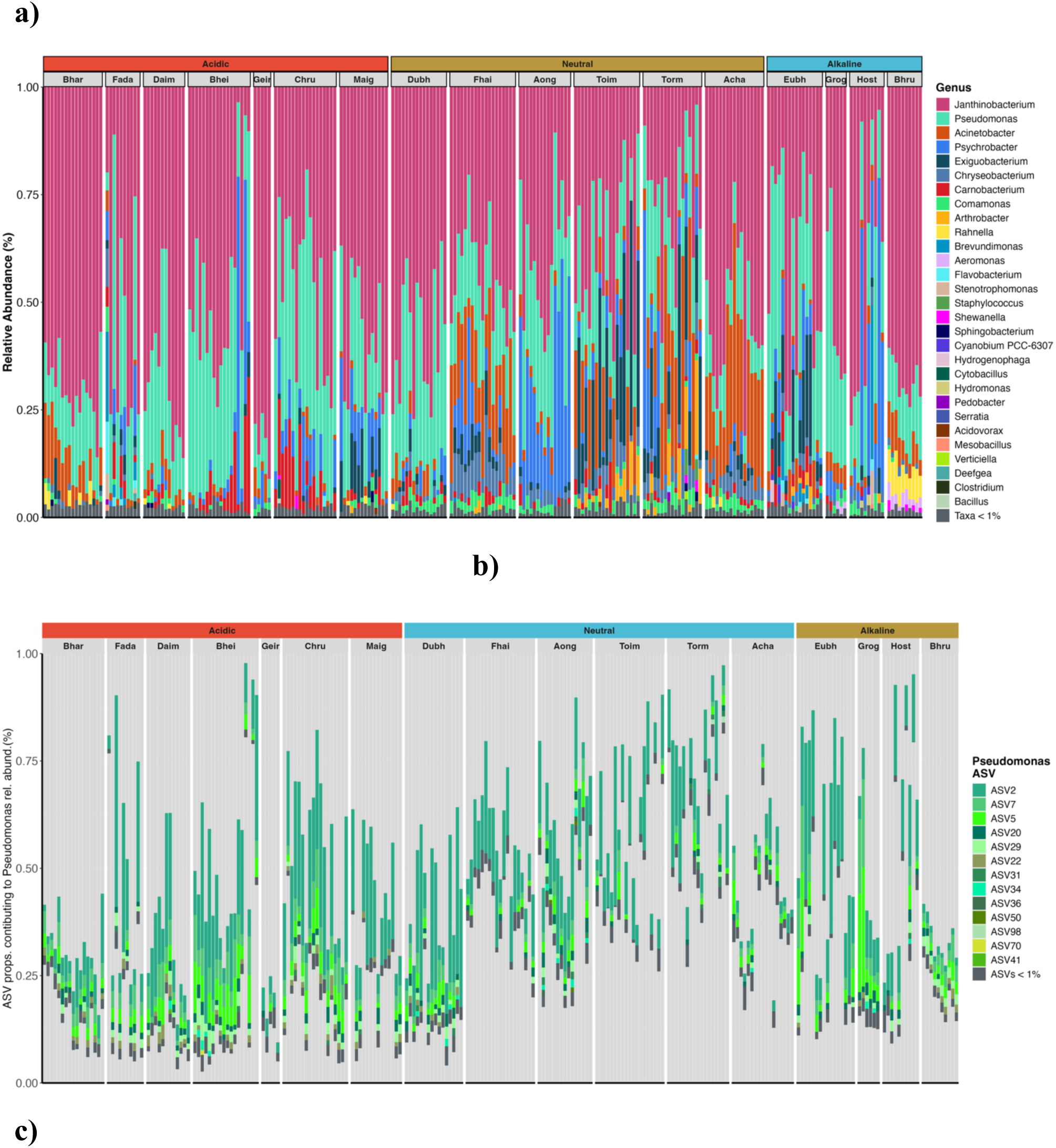
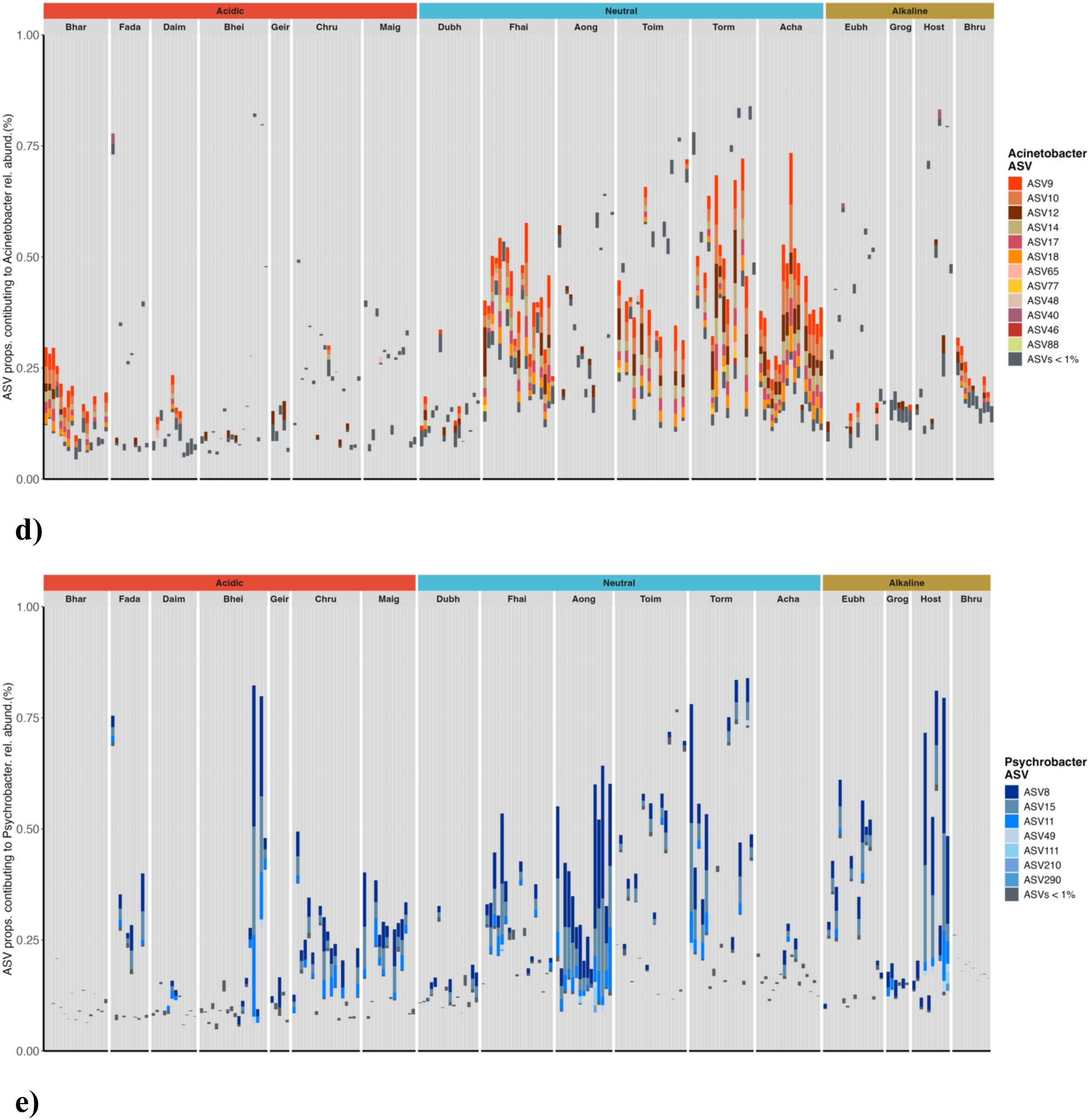
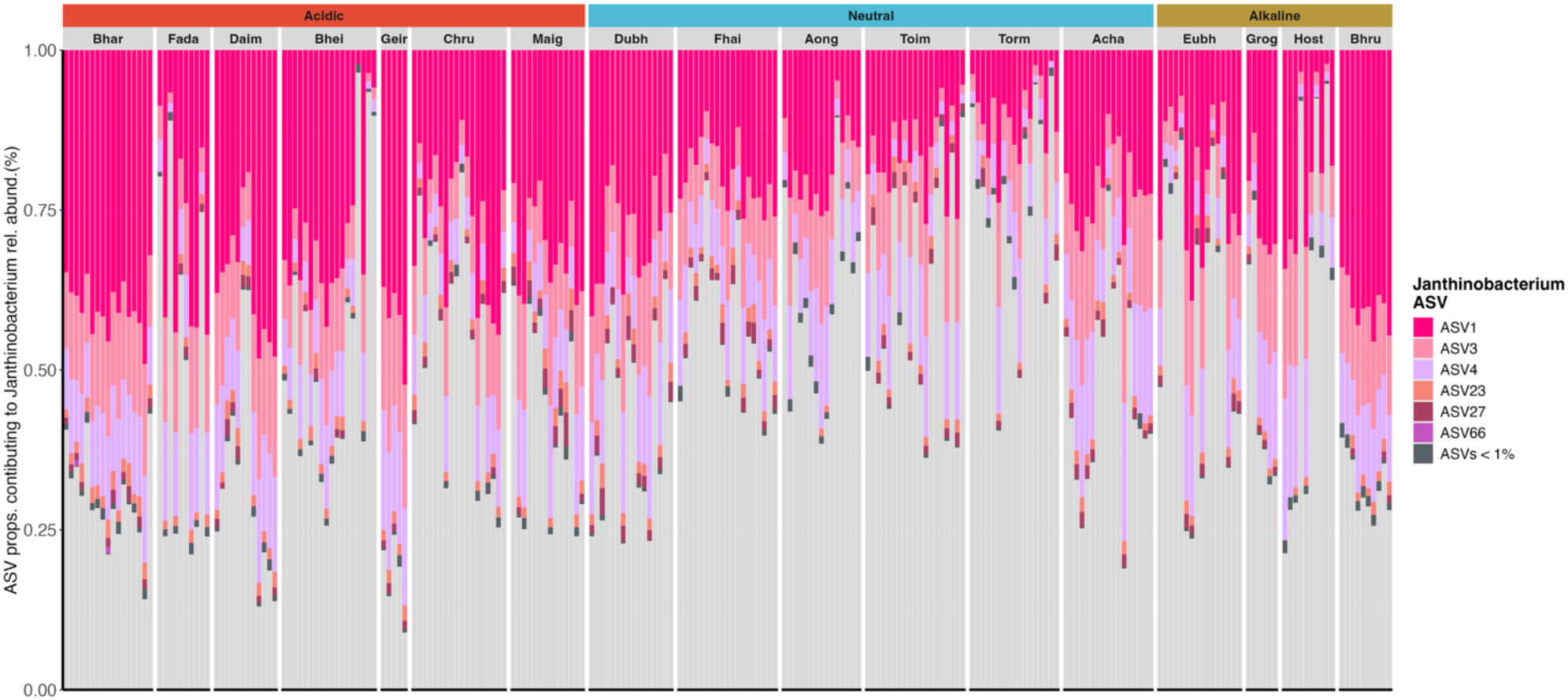
The skin microbiome composition of wild three-spined stickleback across 17 North Uist lochs. **a)** Taxonomic summary of the skin microbiome of each individual stickleback fish sample. Taxa are displayed at the genus-level of classification; any taxa present at <1% mean relative abundance is grouped into “Taxa <1%” and shown in grey. Samples are grouped by their pH classification: acidic (**red**), neutral (**gold**), and alkaline (**blue**), and by sampling location (loch/lake). Each vertical bar represents the skin microbiome of an individual fish (n = 239), with colours corresponding to genera ranked by the highest mean relative abundance. **b–e)** Within-genus ASV composition for the four most abundant genera: **b)** *Pseudomonas*, **c)** *Acinetobacter*, **d)** *Psychrobacter*, **e)** *Janthinobacterium*. Each bar represents an individual fish, and colours indicate distinct ASVs within each genus, as shown in the corresponding legend. Any ASV not within the focal genus is coloured grey in addition to any ASVs with relative abundances <1%.

### Host origin and environmental pH shape microbiome composition and diversity on the stickleback skin

We compared alpha diversity of the stickleback skin microbiome from individuals across lochs to evaluate variation associated with host origin. Both Shannon (Kruskal–Wallis χ²(16) = 140.77, *p* < 0.001) and Simpson (Kruskal–Wallis χ² (16) = 148.42, *p* < 0.001) diversity differed significantly among lochs. Lochs such as Fhai, Toim, Torm, and Acha exhibited higher diversity, whereas Geir showed lower diversity (**Figure 2a; Supplementary Figure 2a**). When grouped by pH classification, Shannon diversity differed significantly among categories (Kruskal–Wallis χ²(2) = 96.68, *p* < 0.001; **Figure 2b**). After accounting for loch as a random effect, stickleback from neutral lochs exhibited higher bacterial diversity than those from acidic lochs (lmer, *β* = 0.50 ± 0.10 SE, *p* < 0.001). Meanwhile, fish from alkaline lochs showed increased diversity relative to acidic lochs, but this difference was not statistically significant (lmer, *β* = 0.22 ± 0.11 SE, *p* = 0.06; **Figure 2b**). Pairwise comparisons further showed that bacterial diversity did not differ significantly between neutral and alkaline lochs (estimate = 0.28 ± 0.12 SE, t = 2.48, *p* = 0.064). Alpha diversity differed significantly between sexes, with females exhibiting higher diversity than males (Welch *t-*test: *t₉₃.₀₇* = 3.07, *p* = 0.003; females = 2.55, males = 2.39; **Figure 2c**). These results indicate that both pH and fish sex independently influence microbial alpha diversity on the stickleback skin.

**Figure 2:**
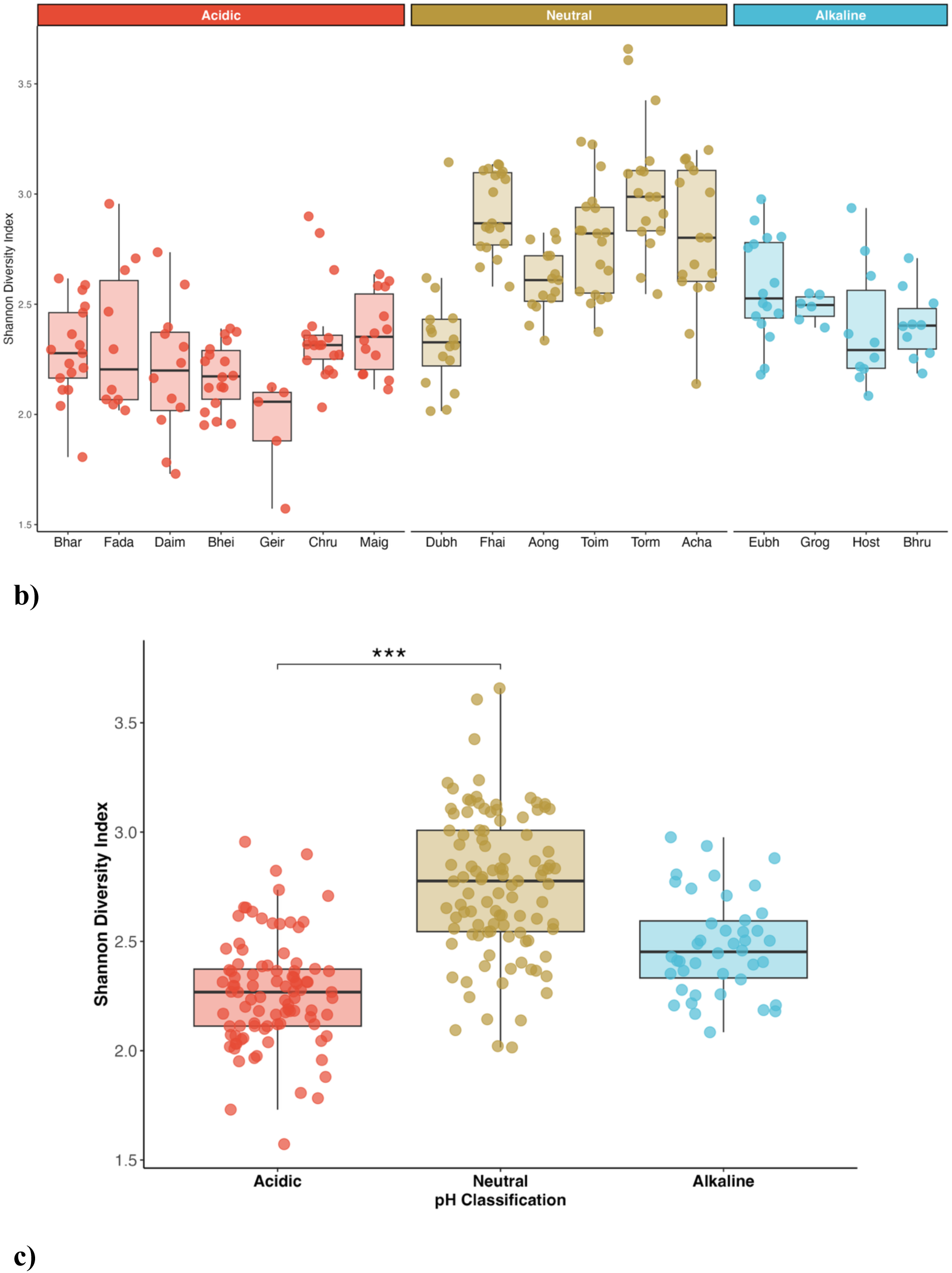
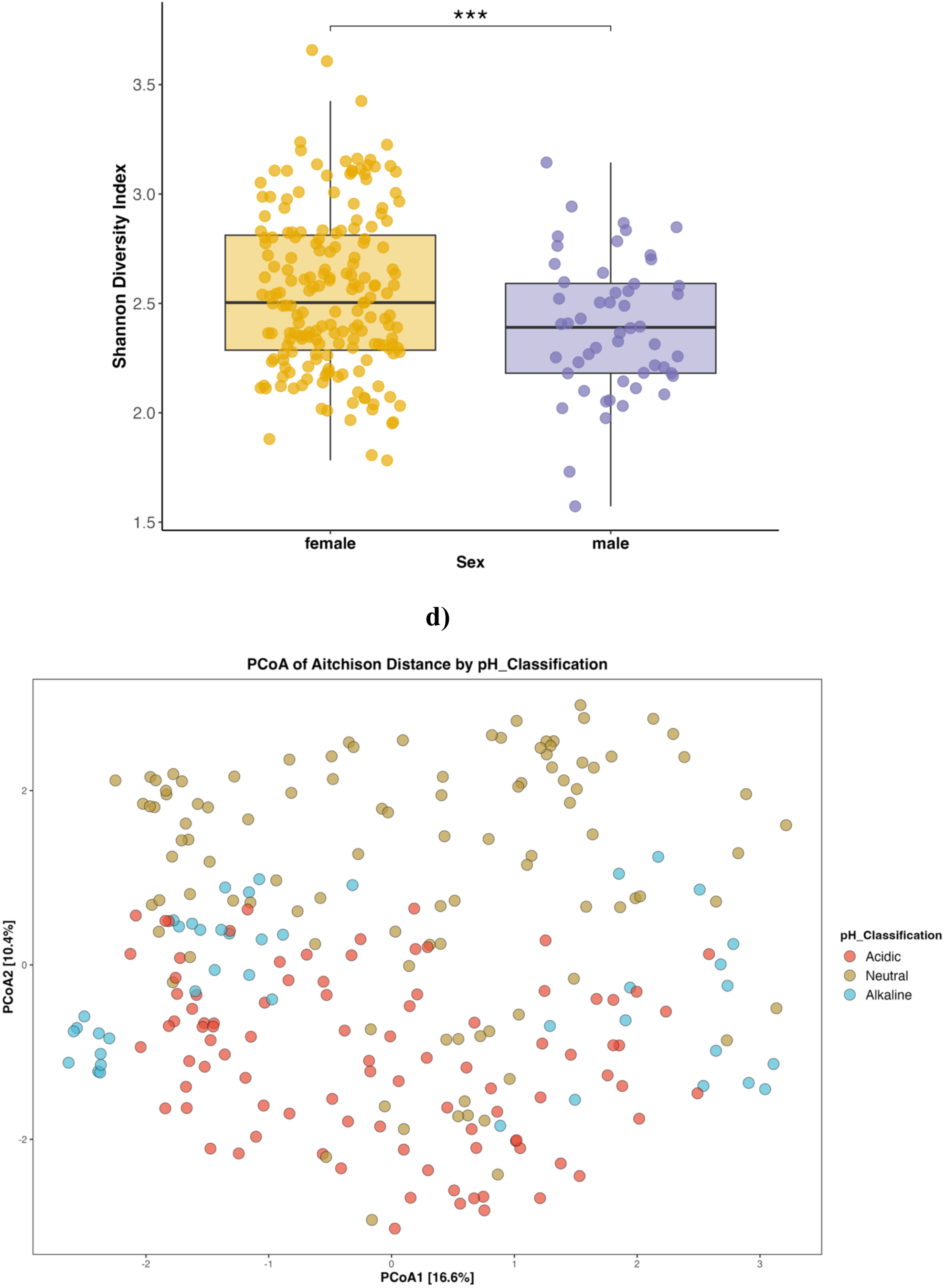
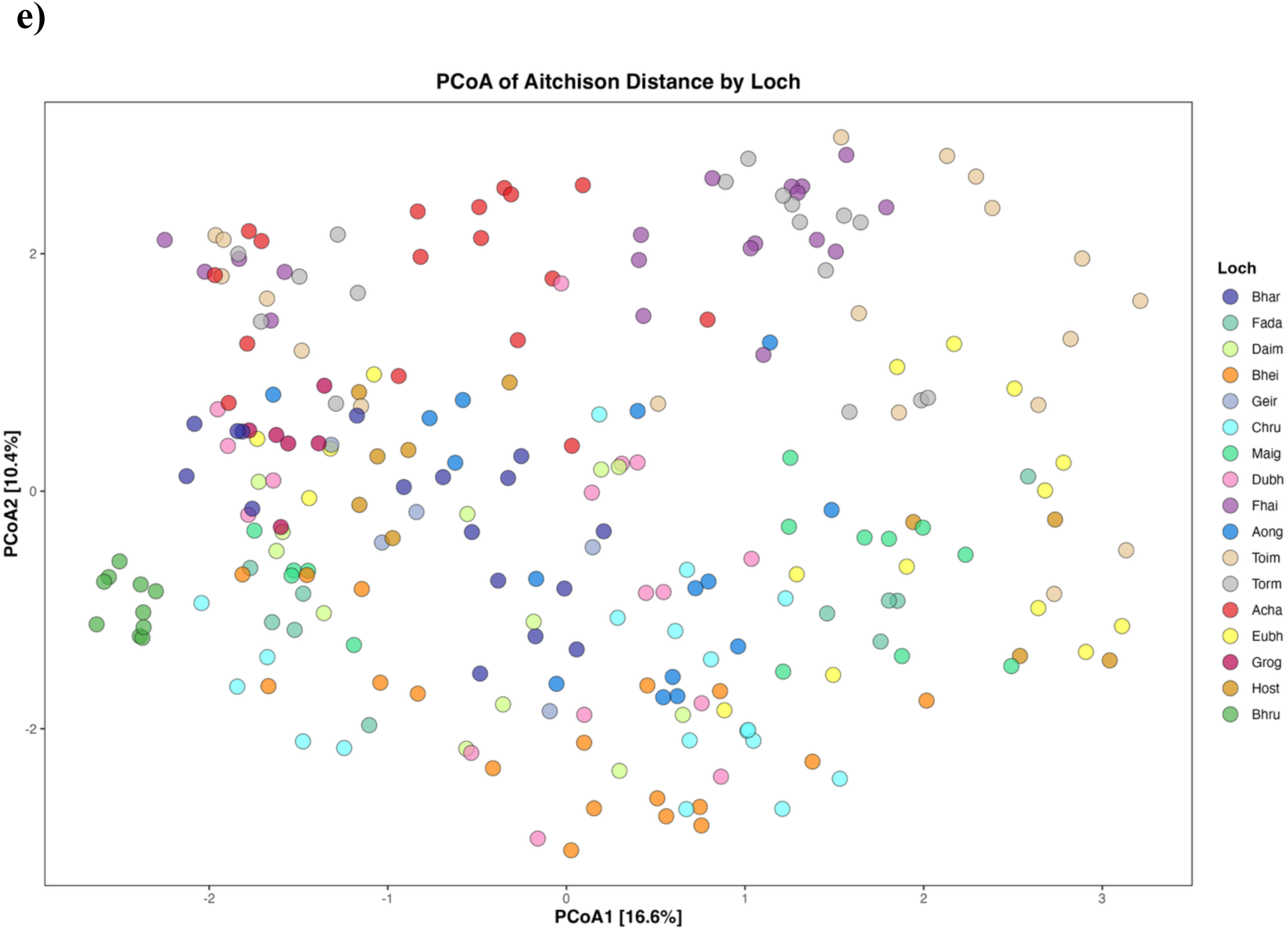
The diversity and community composition of the stickleback skin microbiome vary with pH and host habitat. **a)** Alpha (Shannon) diversity differs significantly between lochs (Kruskal–Wallis χ²(16) = 140.77, *p* < 0.001). Boxplot colours correspond to pH as shown on the x-axis. **b)** Shannon diversity grouped by pH classification (acidic, alkaline, neutral), with sticklebacks from neutral pH lochs hosting a more diverse bacteria (Kruskal–Wallis χ²(2) = 96.68, *p* < 0.001). **c)** Alpha diversity differed between male and female sticklebacks, with females having a higher microbial diversity on average compared to males (p = 0.003). **d, e)** Community-wide microbiome composition changes significantly by pH classification (PERMANOVA: R² = 0.07, F = 8.47, *p* < 0.001) and loch habitat (R² = 0.27, F = 5.04, P < 0.001). pH classification is coloured as acidic (red), neutral (gold), and alkaline (blue), and all lochs are given unique colours.

Similarly, beta diversity analysis using Aitchison and Bray-Curtis distances revealed compositional differences in the stickleback skin microbiome across loch and pH classifications (**Figure 2d,e; Supplementary Figure 2b,c**). The first two axes explained 16.6% and 10.4% of the total variance, respectively (**Figure 2d,e**). When samples were grouped by pH classification, PERMANOVA indicated significant differences in community composition (Aitchison: R² = 0.07, F = 8.47, *p* < 0.001; Bray–Curtis: R² = 0.10, F = 13.18, P < 0.001), indicating a significant interaction between pH and skin microbiome composition. (**Figure 2d**).

The skin microbiome composition of stickleback was signficiantly associated with the originating loch habitat. When samples were coloured by host identity (i.e., loch) (PERMANOVA: Aitchison R² = 0.27, F = 5.04, P < 0.001; Bray–Curtis R² = 0.39, F = 8.98, P < 0.001), points were broadly dispersed throughout the ordination space with considerable overlap between lochs, indicating within-loch variability and limited discrete clustering by individual loch. Nevertheless, some lochs, particularly Bhru (green), formed distinct clusters separate from other lochs with similar pH classifications.

### Influence of host and environmental factors on stickleback skin microbiome composition

Metal concentrations varied across lochs and exhibited clear site-specific patterns (**Supplementary Figure 3**). Zinc concentrations ranged from 14.8 to 45.1 µg/L, with all lochs exceeding the environmental quality standards (EQS) for freshwater set by the UK Technical Advisory Group (UKTAG) and the Scottish Environment Protection Agency (SEPA), 10.9 µg/L (SEPA, 2020; UKTAG, 2014). The highest concentrations of zinc were observed in Grog, Fada, and Maig, surpassing 30 µg/L (**Supplementary Figure 3**). Copper concentrations ranged from 1.7 to 9.8 µg/L, with every loch exceeding the UK guideline of 1 µg/L. The highest copper level was recorded in Toim (9.83 µg/L). In contrast, lead concentrations were comparatively low, ranging from 0.07 to 0.52 µg/L, and remained below the UKTAG/SEPA freshwater standard of 1.2 µg/L at all sites (**Supplementary Figure 3**).

Accordingly, we tested associations among host traits, environmental variables, and the microbiome using PERMANOVA models based on both Bray-Curtis and Aitchison distance metrics to identify the key factors shaping microbiome composition. Both loch and sampling date were significantly associated with differences in microbiome composition (PERMANOVA, *P* = 0.001, **Table 2**). While sampling date and loch were correlated due to field sampling constraints, loch remained a significant predictor of microbiome composition when sampling date was included in the model (**Table 2**). Sampling date accounted for 17.9% (Bray–Curtis) and 12.5% (Aitchison) of the variation, while loch explained an additional 10.9% and 7.4%, respectively (**Table 2**).

**Table 2:** Association between metadata variables and the composition of the microbiome.

| **Data** | Bray-Curtis distance<br>(p-value/( R<sup>2</sup>-value)) | Aitchison distance<br>(p-value/ (R<sup>2</sup>-value)) |
| --- | --- | --- |
| **Sampling date** | 0.001 (0.179) | 0.001 (0.125) |
| **Sex** | 0.004 (0.027) | 0.001 (0.010) |
| **Length** | 0.024 (0.008) | 0.006 (0.007) |
| **pH_classification** | 0.001 (0.025) | 0.001 (0.016) |
| **Conductivity** | 0.002 (0.015) | 0.010 (0.014) |
| **Copper (Cu)** | 0.001 (0.018) | 0.001 (0.015) |
| **Zinc (Zn)** | 0.014 (0.009) | 0.004 (0.009) |
| **Lead (Pb)** | 0.034 (0.032) | 0.001 (0.012) |
| **Loch** | 0.001 (0.109) | 0.001 (0.074) |

Among environmental parameters, pH classification, conductivity, and metal concentrations (zinc, lead, and copper) emerged as significant drivers of microbiome composition. Specifically, pH classification explained 2.5% of the variance (Bray–Curtis, P = 0.001) and 1.6% (Aitchison, P = 0.001), while Pb explained 1.2% (Bray–Curtis, P = 0.001) and 3.2% (Aitchison, P = 0.001). Conductivity was also significantly associated with microbiome composition under both distance metrics (Bray–Curtis: R² = 0.015, P = 0.002; Aitchison: R² = 0.014, P = 0.014). Cu and Zn were also significantly associated with microbiome composition, although Zn showed the smallest effect size among the three metals.

In contrast to environmental drivers, host-associated traits explained relatively little variance in microbiome composition. Sex was significantly associated with community structure based on the Bray–Curtis distance (R² = 0.027, P = 0.004) but showed a weaker effect with the Aitchison distance (R² = 0.010, P = 0.001). Length exhibited a small but significant association with microbiome composition under both distance metrics (Bray–Curtis: R² = 0.008, P = 0.024; Aitchison: R² = 0.007, P = 0.006). Overall, environmental factors (pH, metals, and conductivity) explained approximately 6.6–10% of the variation in microbiome composition, while host-associated factors (sex, length) accounted for less than 3.5% (**Table 2**). However, sampling day and habitat (loch) remained the dominant sources of variation, together explaining over 25% of the differences in microbiome composition across both distance metrics.

### Differentially abundant taxa associated with pH classification and host sex

Given the contributions of environmental pH classification and host sex to variation in microbiome composition, we next examined whether specific bacterial taxa drove these observations. Differential abundance analysis with DESeq2 identified 27 ASVs that varied significantly across pH categories (**Figure 3a,b; Supplementary Table 4**). Most of these ASVs were relatively abundant in the dataset, with 22 exhibiting a mean relative abundance greater than 0.1%. Based on abundance patterns, taxonomic identity, and associations with other metadata variables, we focused subsequent analyses on four ASVs (ASV6, ASV16, ASV28, and ASV38) that showed pronounced differences across pH classifications.

**Figure 3:**
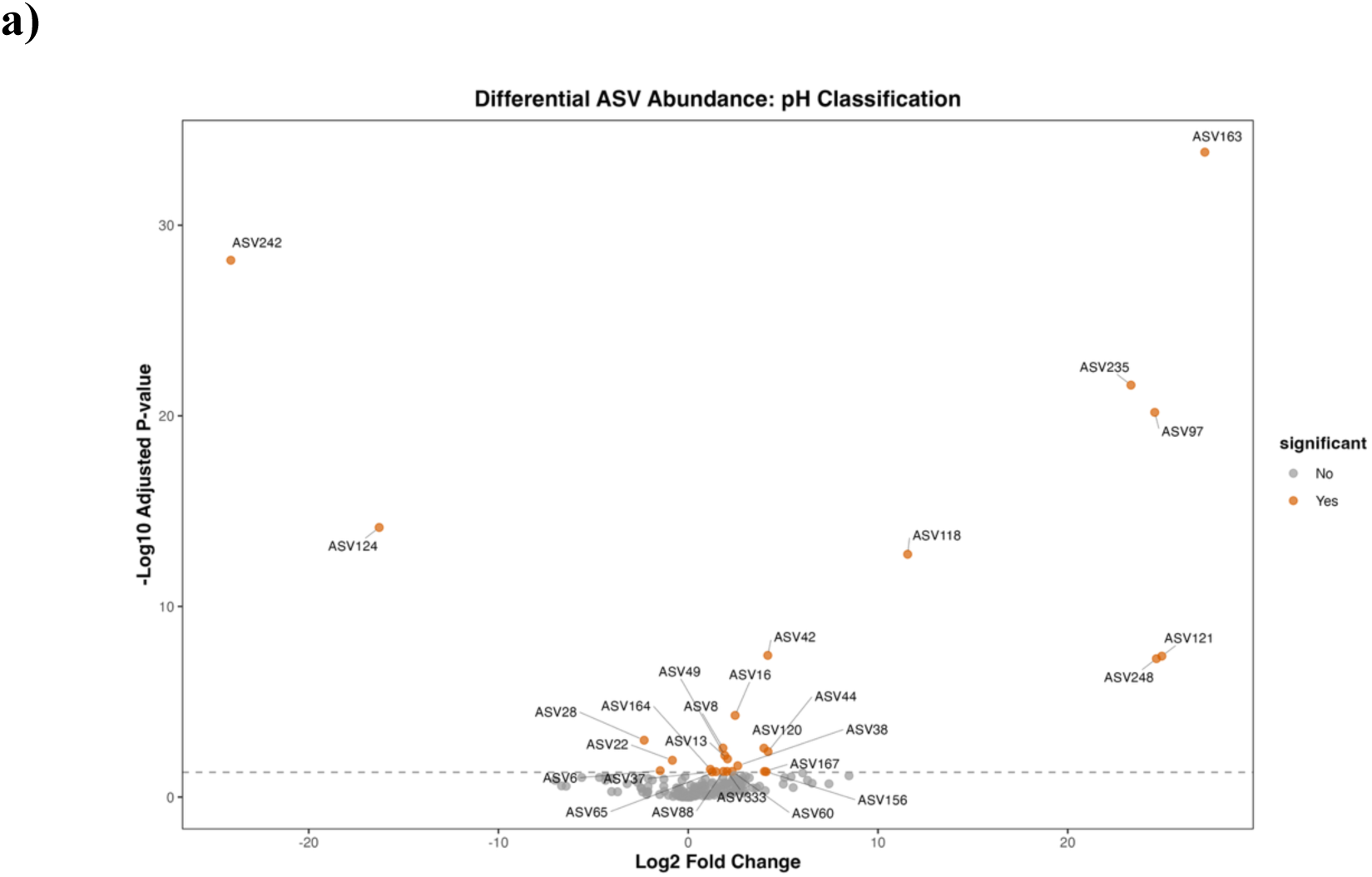
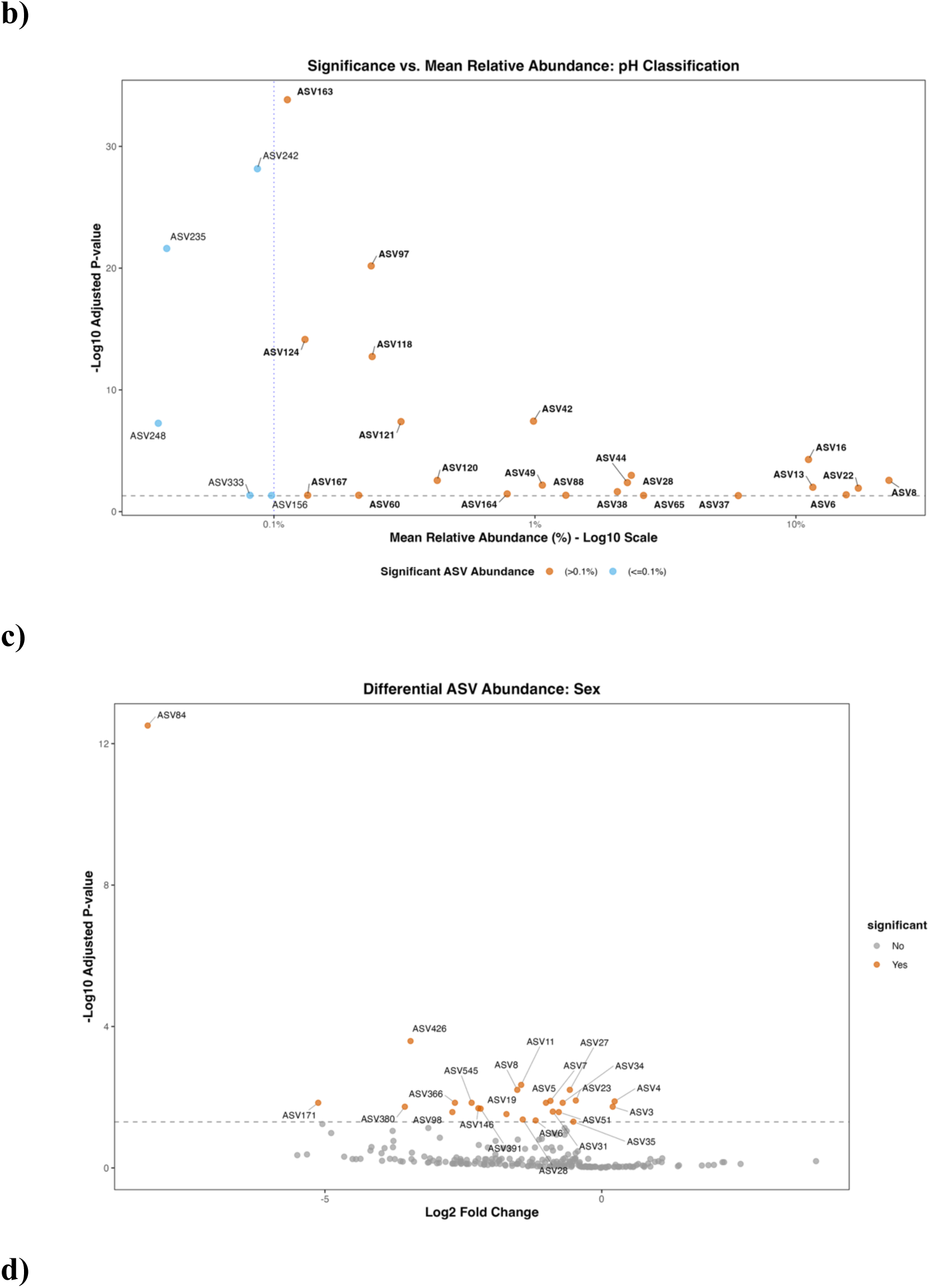
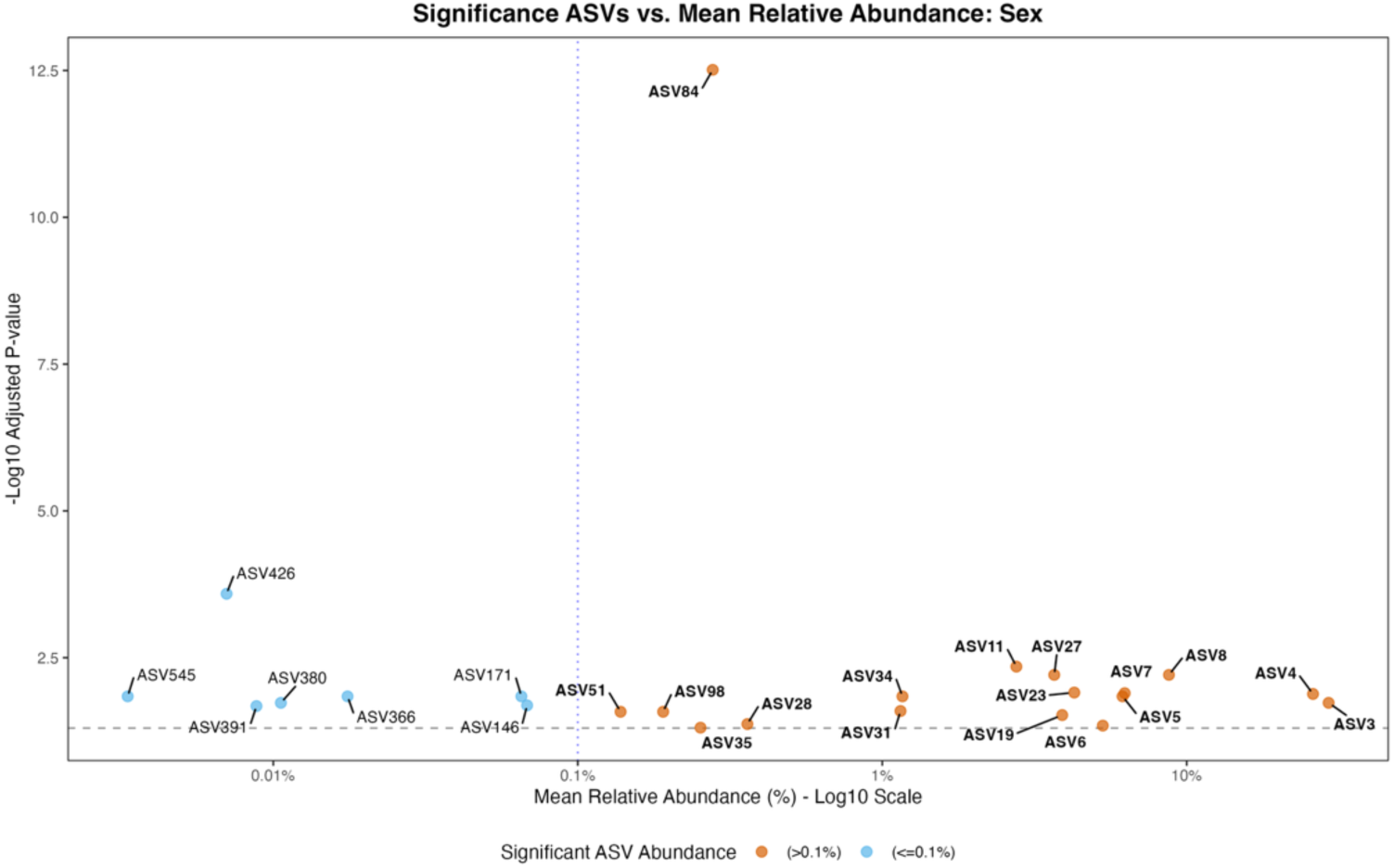
Differentially abundant ASVs associated with pH classification and host sex. (a) Volcano plot showing ASVs differing across pH classifications. (b) Mean relative abundance versus significance for pH-associated ASVs. (c) Volcano plot showing ASVs differing between male and female hosts. (d) Mean relative abundance versus significance for sex-associated ASVs. Points represent individual ASVs. The x-axis shows log₂ fold change (volcano plots) or mean relative abundance (log₁₀ scale), while the y-axis shows the log₁₀ adjusted *p*-value from DESeq2 analysis.

ASV6 (*Carnobacterium*) exhibited its highest relative abundance in fish sampled from acidic lochs, where both the median abundance and variability exceeded those observed in neutral and alkaline environments (**Supplementary Figure 4a–d**). ASV28 (*Sphingobacterium*) showed a similar distribution, with higher median abundance and greater variability in acidic lochs than in other pH classes (**Supplementary Figure 5a–d**). ASV6 had significantly higher relative abundances in females, whereas ASV28 showed no clear separation between sexes in neutral and alkaline lochs but had higher abundances in acidic lochs. Elevated abundances of these taxa were contributed by samples from several acidic lochs, including Fada, Bhei, Chru, Bhar, and Daim, although most individuals across sites exhibited low relative abundances.

In contrast, ASV16 (*Chryseobacterium*) and ASV38 (*Arthrobacter*) were most abundant in neutral lochs. ASV16 displayed the highest relative abundances in neutral environments, with consistently low abundances in acidic systems and intermediate levels in alkaline lochs (**Supplementary Figure 6a–d**). ASV38 exhibited a similar pattern, with the lowest abundances in acidic systems and increased abundance in neutral lochs, with moderate levels in some alkaline samples (**Supplementary Figure 7a–d**). High abundances were observed across both males and females and across multiple sites, including Fhai, Toim, Torm, and Acha.

To further examine host-associated patterns, we performed an equivalent analysis to test for sex-associated taxa. This analysis revealed 24 ASVs that differed significantly between male and female hosts (**Figure 3c,d; Supplementary Table 5**), of which 16 had a mean relative abundance greater than 0.1%. Applying the same criteria of abundance, taxonomic identity, and ecological relevance, we conducted further analyses of two ASVs (ASV6 and ASV8) that showed strong associations with host sex and other metadata variables. ASV6 (*Carnobacterium*) and ASV8 (*Psychrobacter*) exhibited higher relative abundance and greater dispersion in female samples than in male samples (**Supplementary Figure 8a-d,9a-d**). Several female individuals exhibited elevated abundances, exceeding 0.3–0.5% (**Supplementary Figure 8a–d**), whereas most male samples had low relative abundances with a narrower distribution. Samples with higher abundances occurred across multiple pH categories and lochs, including Chru, Fada, Bhei, Aong, Torm, Eubh, and Host.

## Discussion

While previous work on the microbiome of three-spined stickleback has focused primarily on gut-associated communities, the microbial composition of external mucosal surfaces in natural environments has remained underexplored. Our primary goal was to investigate the composition and diversity of these microbial communities and assess the relative influence of environmental factors and host traits on their composition. We found that microbial community composition and diversity were primarily shaped by habitat (loch), pH, metal concentrations, and host-associated traits, including sex or body size. Additionally, we identified a set of dominant and consistently present bacterial genera across habitats, suggesting that the stickleback skin harbours a core microbiota with potential ecological or symbiotic roles.

### The stickleback skin microbiome showed a conserved and dominant bacterial genus

Across all individuals and lochs, the skin microbiome was dominated by genera including *Janthinobacterium, Pseudomonas, Acinetobacter*, and *Psychrobacter*. These taxa were consistently abundant, with *Pseudomonas* exhibiting the highest within-genus ASV-diversity. While the dominance of *Janthinobacterium* appears unique to this study, the bacterial genera identified in this study are consistent with findings from other fish species, where they have been reported as common and functionally important members of the skin microbiota (Krotman et al., 2020; Larsen et al., 2013; Sultana et al., 2022; Zou et al., 2023). The consistent presence of *Janthinobacterium* as a core taxon on stickleback skin suggests that this genus may play an important role in host defence or microbiome stability. *Janthinobacterium* spp. are known for producing violacein, a pigment with potent antimicrobial and antifungal properties that may contribute to host defence by suppressing opportunistic or pathogenic microbes (Asencio et al., 2014; Baricz et al., 2018; Brucker et al., 2008; Lorentsen et al., 2025). Similarly, *Pseudomonas* spp., which are metabolically versatile and commonly commensal in aquatic animals, may contribute to biofilm formation, immune modulation, and pollutant detoxification, as reported in other fish species (Galina et al., 2015; Sultana et al., 2022). Their widespread occurrence and variability across sites may reflect strong environmental plasticity, suggesting that these bacteria could play flexible symbiotic roles on the host skin under differing environmental conditions.

*Psychrobacter*, a cold-tolerant genus frequently found in aquatic and polar environments (Bakermans, 2018; Rodrigues et al., 2009), was also consistently represented across individuals. Although its role in the fish skin microbiome is not well-characterised, variation among its ASVs suggests the presence of closely related strains that may be part of the resident mucosal microbiota. These taxa could contribute to community stability under fluctuating environmental conditions, potentially supporting microbiome homeostasis in cold environments or in waters with elevated metal concentrations. (Chiarello et al., 2017; Lowrey et al., 2015; Muñoz-Villagrán et al., 2018; Sehnal et al., 2021b). While often associated with opportunistic infections in clinical settings (Fahy et al., 2023; Howard et al., 2012), *Acinetobacter* is also widely distributed in aquatic environments, where it contributes to nitrogen metabolism and the degradation of pollutants (Jung & Park, 2015; Mao et al., 2024; Touchon et al., 2014). The high ASV-level diversity observed within *Acinetobacter* across lochs may reflect the presence of multiple closely related strains adapted to local environmental conditions. Such variation could reflect adaptation to factors such as metal exposure or other chemical gradients, highlighting the potential ecological flexibility of this genus in fish microbiomes (Jung & Park, 2015; Zhang et al., 2024). Collectively, these taxa likely represent a functional core microbiome contributing to mucosal homeostasis, colonisation resistance, and resilience to environmental stressors in stickleback. Nonetheless, metagenomic and functional approaches will be necessary to confirm the metabolic capabilities and ecological roles of these taxa.

### Environmental factors are a major influence on microbiome composition

Our analyses revealed that environmental conditions were key drivers of microbiome composition. Metal concentrations (Zn, Cu, Pb), pH, and conductivity showed consistent effects across both beta diversity distance metrics. These findings support the growing evidence that abiotic variables function as ecological filters, shaping mucosal microbial communities and influencing the structure of host-associated microbiomes (Lokesh & Kiron, 2016; F.-É. Sylvain et al., 2020). Fish from neutral and alkaline lochs had significantly higher alpha diversity than those from acidic environments, suggesting that microbial richness increases under less physiologically stressful conditions (Lauber Christian et al., 2009; Sylvain et al., 2016). pH can influence microbial membrane stability, nutrient uptake, and enzymatic activity, thereby constraining the taxa able to persist under different chemical conditions (Delgado-Baquerizo et al., 2018; Lauber Christian et al., 2009). Accordingly, pH may act as an environmental filter, favouring acid-tolerant bacteria in low-pH lochs and alkaline-tolerant taxa in alkaline conditions, which, in turn, shape the overall microbiome composition, as reflected by differences in beta-diversity (Atasoy et al., 2024; Jin & Kirk, 2018). Furthermore, the distinct separation of Loch Bhru from the other alkaline lochs may reflect location-specific environmental conditions. Despite having a similar pH, its geographic isolation (**Figure 1B**) may expose it to different catchment characteristics, hydrological changes, or other unmeasured environmental factors that influence microbiome communities.

Heavy metals, particularly Cu, Zn, and Pb, were significant predictors of microbiome composition across both beta-diversity metrics. These metals are known to disrupt microbial homeostasis by selectively inhibiting sensitive taxa while enabling the persistence of metal-resistant or metabolically versatile genera (Mohammed & Arias, 2015; Qian et al., 2020). Copper can be highly toxic to microorganisms at elevated concentrations and may induce copper-resistance mechanisms (Trevors & Cotter, 1990). In the UK, metals such as Cu, Zn, and Pb are recognised environmental risk factors due to their accumulation in aquatic organisms and ecosystems (Donnachie et al., 2014; JNCC, 2023). Our findings therefore support growing evidence that bioavailable metals in freshwater systems can act as important drivers of mucosal microbiome composition (Flemming & Trevors, 1989), although further work is needed to elucidate direct associations between metal exposure and specific microbial taxa or functional responses. Conductivity, a less frequently discussed driver in fish microbiome studies, also significantly influenced the microbiome composition. As a proxy for ion concentration and overall salinity, conductivity may influence microbial adhesion, nutrient diffusion, and osmotic regulation within skin-associated microbial communities (A. G. Bell et al., 2024; Krotman et al., 2020). Similar findings have been reported in other fish species, where skin microbiome beta diversity varies in response to environmental factors such as temperature, conductivity, and pH (Krotman et al., 2020; Sehnal et al., 2021b).

### Effects of host traits on microbiome composition

In contrast to the pronounced influence of environmental factors, host-related traits exhibited comparatively minor effects on microbiome composition. Sex showed a statistically significant association with alpha diversity, with female fish displaying higher Shannon diversity and this pattern was consistent across beta diversity measures. These findings align with prior studies (Bolnick et al., 2014; Elisa Casadei et al., 2023; E. Casadei et al., 2023) showing that host sex can influence microbial community composition. These differences reflect sex-specific physiological or behavioural differences during the breeding season. In sticklebacks, males engage in energetically costly nest building and parental care, which may alter immune function and microbial colonisation dynamics (Álvarez-Quintero et al., 2021). Fish length showed a marginal association with microbial composition across both distance metrics, which could reflect the relatively uniform size of sticklebacks sampled across freshwater lochs. The limited variation in standard length among individuals may have reduced the power to detect strong length-related effects on microbial community composition.

### Host habitat and temporal effects on the microbiome

While specific environmental parameters were strong predictors of microbiome composition, habitat identity (loch) and temporal effects (sampling date) explained additional variation, highlighting the importance of these factors in shaping microbial communities (Sadeghi et al., 2023; F. Sylvain et al., 2020). This pattern suggests that the external microbiome responds dynamically to short-term environmental fluctuations such as rainfall, runoff, changes in water chemistry, or shifts in host immune status (Minich et al., 2020). These findings further emphasize the importance of temporal replication in fish microbiome studies and suggest that surface-associated microbiota may serve as sensitive indicators of environmental instability (Daniela Rosado et al., 2021). Nevertheless, these effects should be interpreted cautiously, as loch identity and sampling day are partially confounded by the sampling design, making it difficult to isolate their independent contributions.

### Environmental and host factors shape the abundance of specific taxa in the stickleback microbiome

Differential abundance analysis identified multiple ASVs associated with pH classification and host sex, indicating that environmental conditions and host-associated factors jointly shape the microbiome composition. Similar interactions between abiotic conditions and host traits have been observed in other fish microbiome studies, where environmental variables structure microbial assemblages, while host characteristics influence the abundance of specific taxa (Ashley G. Bell et al., 2024; Boutin et al., 2014). Two taxa, *Carnobacterium* (ASV6) and *Sphingobacterium* (ASV28), were associated with fish from acidic lochs, implying that acidic environments may favour colonisation or persistence of these taxa on stickleback skin.

*Carnobacterium* are lactic acid bacteria capable of fermentative metabolism and the production of organic acids and antimicrobial compounds (e.g., bacteriocins) (González-Gragera et al., 2024; Ringø, 2024). Accordingly, members of this genus are reported as commensals or probiotics in fish microbiomes, contributing to community stability and host defence by suppressing opportunistic pathogens and modulating local microbial environments (Menanteau-Ledouble et al., 2022). These physiological traits provide a competitive advantage and may help explain their increased abundance in acidic environments where bacterial communities must tolerate low pH and fluctuating nutrient conditions (Tang et al., 2023). Similarly, *Sphingobacterium* is widely distributed in freshwater and soil environments and is commonly associated with the degradation of complex organic compounds (Kakumanu et al., 2021; Luan et al., 2023). Species within this genus possess diverse metabolic capabilities, including the breakdown of polysaccharides and other high-molecular-weight organic substrates commonly found in freshwater ecosystems (Kim Sung & Wang Sharon, 2024; Song et al., 2020). Acidic freshwater systems frequently receive substantial inputs of humic and fulvic acids due to runoff from surrounding soils, resulting in elevated concentrations of complex organic matter (Moody, 2024; Oliver et al., 1983). The increased abundance of *Sphingobacterium* in fish from acidic lochs may therefore reflect adaptation to organic-rich, low-pH environments, where their metabolic capabilities enable them to exploit abundant substrates in these systems.

In contrast, *Chryseobacterium* and *Arthrobacter* were more abundant in neutral lochs. Species of Chryseobacterium are frequently reported from fish skin, gills, and aquaculture systems and function as commensal members of the fish-associated microbiota (Wang et al., 2025). Similarly, *Arthrobacter* species are widely distributed freshwater and soil bacteria known for their metabolic versatility and ability to degrade diverse organic compounds (Jones & Keddie, 2006). Both genera typically grow optimally at near-neutral pH and exhibit reduced growth or activity under strongly acidic conditions(Ilardi et al., 2009; Jones & Keddie, 2006; Kim et al., 2008; Margesin et al., 2004). Their enrichment on stickleback skin in neutral lochs, therefore, likely reflects physiological preferences for moderate pH conditions, where environmental pH is closer to their growth optimum. In addition to pH-associated taxa, several ASVs were associated with host sex. Both *Carnobacterium* (ASV6) and *Psychrobacter* (ASV8) were more abundant in females than in males. Sex-associated differences in microbiomes have been reported across vertebrates and are often linked to hormonal regulation of immunity and mucosal surfaces (Bates et al., 2023; Elderman et al., 2018; Fransen et al., 2017). Although these ASVs have not been linked with sex-specific variation in fish microbiomes, their higher abundance and dispersion among females across multiple lochs may reflect differences in skin mucus composition, immune activity, or breeding behaviour, given that sampling occurred during the reproductive period (Sundaray et al., 2025).

While our results identify several taxa associated with pH and host sex, the mechanisms behind these patterns remain unclear. Future research involving functional genomic analyses and experimental manipulation of pH levels would help determine whether these taxa are directly affected by environmental conditions or indirectly influenced through host physiological responses. Additionally, longitudinal sampling across seasons and life stages (or using a gnotobiotic stickleback) could show whether sex-associated ASV patterns remain consistent over time or change with environmental factors and host reproductive cycles.

### Ecological implications and significance

These findings have significant ecological implications. Detecting microbiome shifts in response to sub-regulatory concentrations of heavy metals suggests that skin-associated microbial communities may be a key indicator for early detection of freshwater pollution (Callewaert et al., 2020; Griffero et al., 2024). Also, identifying a stable core microbiota supports the idea that key bacterial taxa contribute to host resilience, potentially maintaining mucosal homeostasis or providing additional functional benefits under environmental stress. Furthermore, the differential abundance of specific ASVs across pH classification suggests that environmental pH could act as an ecological filter, favouring bacterial taxa with physiological traits suited to particular chemical conditions. While the sex-associated ASVs suggest that host biological traits can modulate the abundance of certain microbes. Finally, this study contributes to the broader understanding of host–microbe–environmental interactions in wild aquatic vertebrates. It reinforces that external microbiomes are not passive reflections of ambient conditions but are dynamically shaped by environmental conditions and microbial adaptation.

### Conclusions

This study provides a comprehensive characterisation of the skin microbiome in wild three-spined sticklebacks. Microbial community structure was predominantly shaped by environmental factors, particularly pH and metal concentrations, as well as host-associated traits. Dominant genera such as *Janthinobacterium, Pseudomonas, Acinetobacter*, and *Psychrobacter* formed a potentially functional core microbiota that was consistent across sites but responsive to local environmental gradients. These results contribute to a growing body of work demonstrating that interactions between environmental variations and host biological traits shape host-associated microbiome composition. Understanding these interactions is critical for predicting how microbiomes may respond to environmental change, particularly in freshwater ecosystems where anthropogenic activities (such as agricultural runoff, mining, and wastewater discharge) increasingly influence pH and water chemistry.

