## Supplemental Data for "The effect of environmental variation on the diversity and composition of the three-spined stickleback microbiome"

### Supplemental Material

**a)**

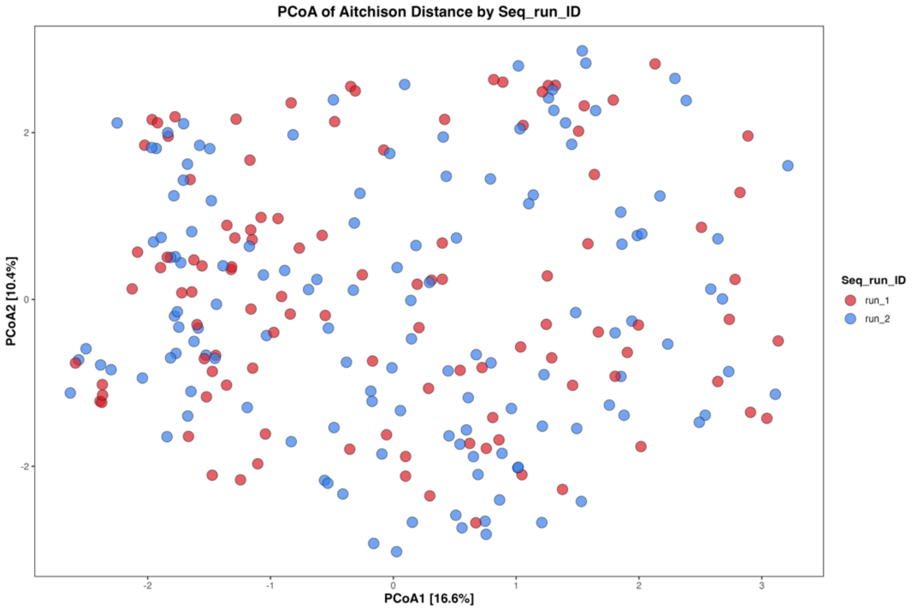


**b)**

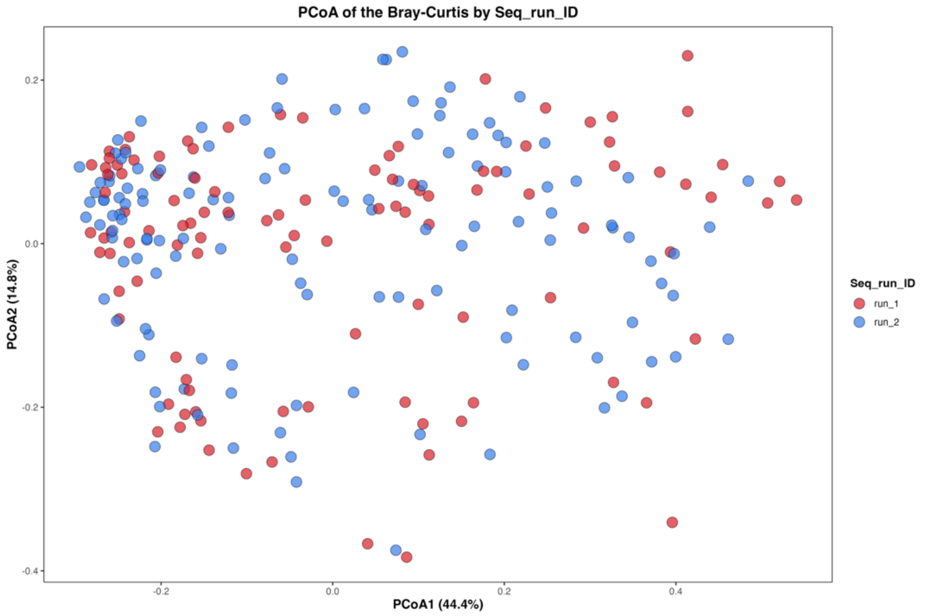


**Supplementary Figure 1:** Assessment of sequencing run effects on microbiome composition. Principal coordinate analysis (PCoA) of microbiome composition based on **(a)** Aitchison distance metrics (PERMANOVA: $R^2 = 0.003$, $F = 0.82$, $P = 0.684$) and **(b)** Bray–Curtis (PERMANOVA: $R^2 = 0.002$, $F = 0.45$, $P = 0.824$). Points represent individual samples and are coloured by sequencing run (Run 1: red, DS1391; Run 2: blue, DS1637). Samples from both runs are broadly intermixed in ordination space, indicating no detectable batch effects.

**a)**

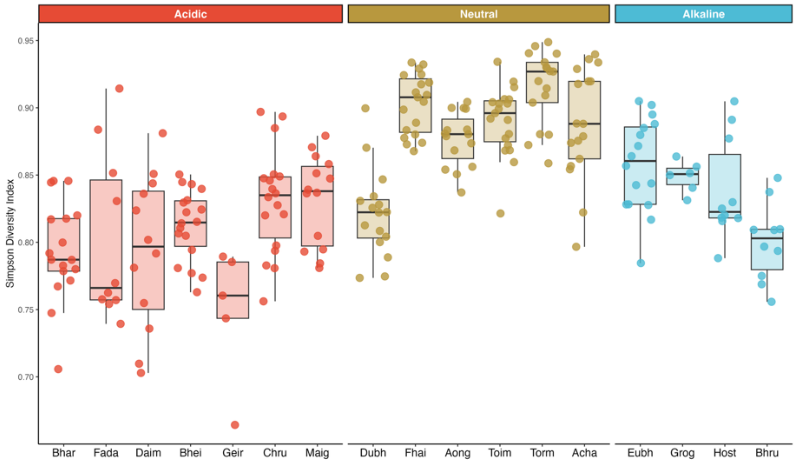


**b)**

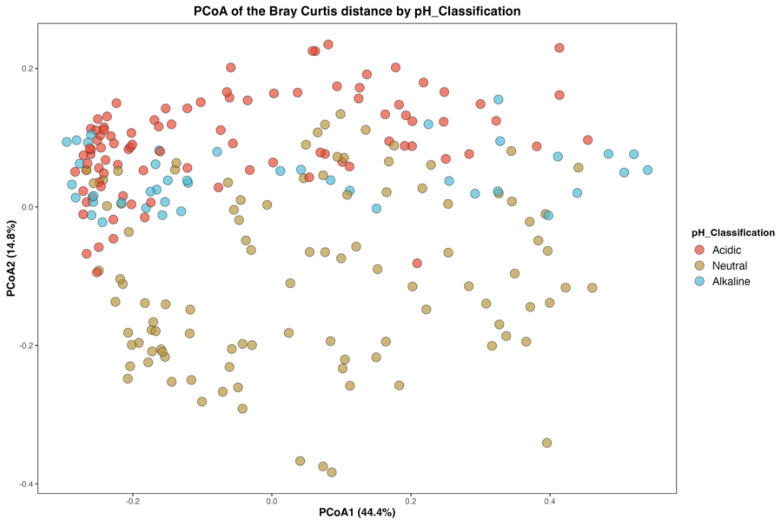


**c)**

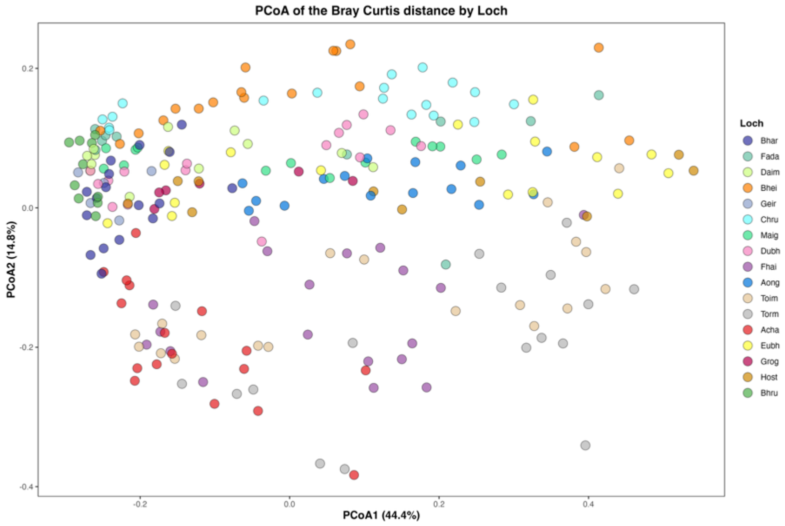


**Supplementary Figure 2:** The diversity and community composition of the stickleback skin microbiome vary with pH and host habitat. **a)** Alpha diversity ((Simpson) differs significantly between lochs (Kruskal–Wallis $\chi^2$ (16)= 148.42, $p$ < 0.001). Boxplot colours correspond to lochs as shown on the x-axis. **b, c)** Community-wide microbiome composition changes significantly among pH classification (PERMANOVA: $R^2 = 0.10$, $F = 13.18$, $P < 0.001$), with Bray-Curtis distances and lochs ($R^2 = 0.39$, $F = 8.98$, $P < 0.001$) respectively. pH classification is coloured as acidic (red), neutral (gold), and alkaline (blue), and all lochs are given unique colours.

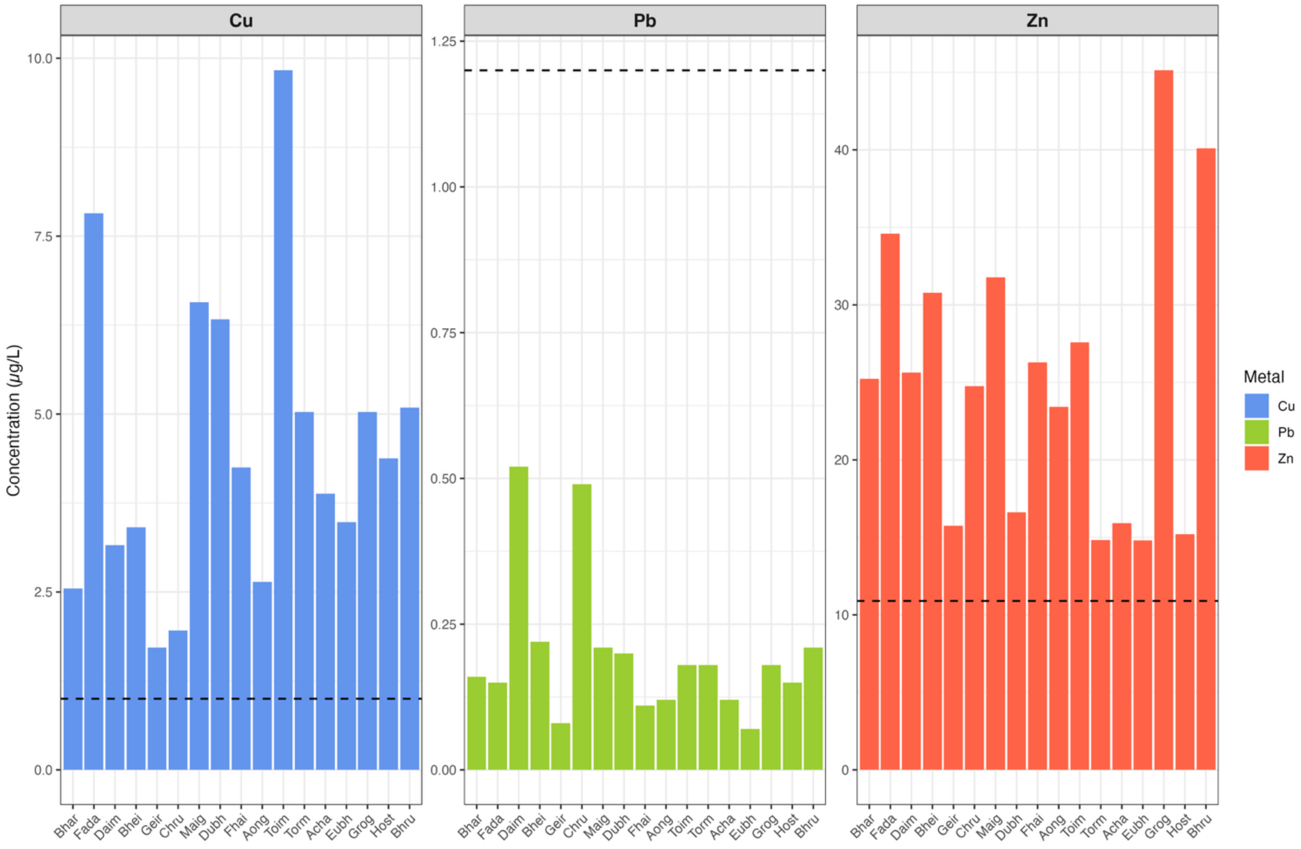


**Supplementary Figure 3:** Concentrations of dissolved trace metals across sampled lochs. Concentrations of copper (Cu), lead (Pb), and zinc (Zn) were measured in water samples from each sampled loch. Bars represent mean metal concentrations (µg $L^{-1}$) for individual lochs. The dashed horizontal line indicates the Environmental Quality Standard (EQS) UKTAG/SEPA threshold for each metal (Cu = 1 µg $L^{-1}$, Zn = 10.9 µg $L^{-1}$, Pb = 1.2 µg $L^{-1}$). Cu and Zn concentrations exceeded EQS thresholds in multiple lochs, whereas Pb concentrations remained below the EQS across all sites.

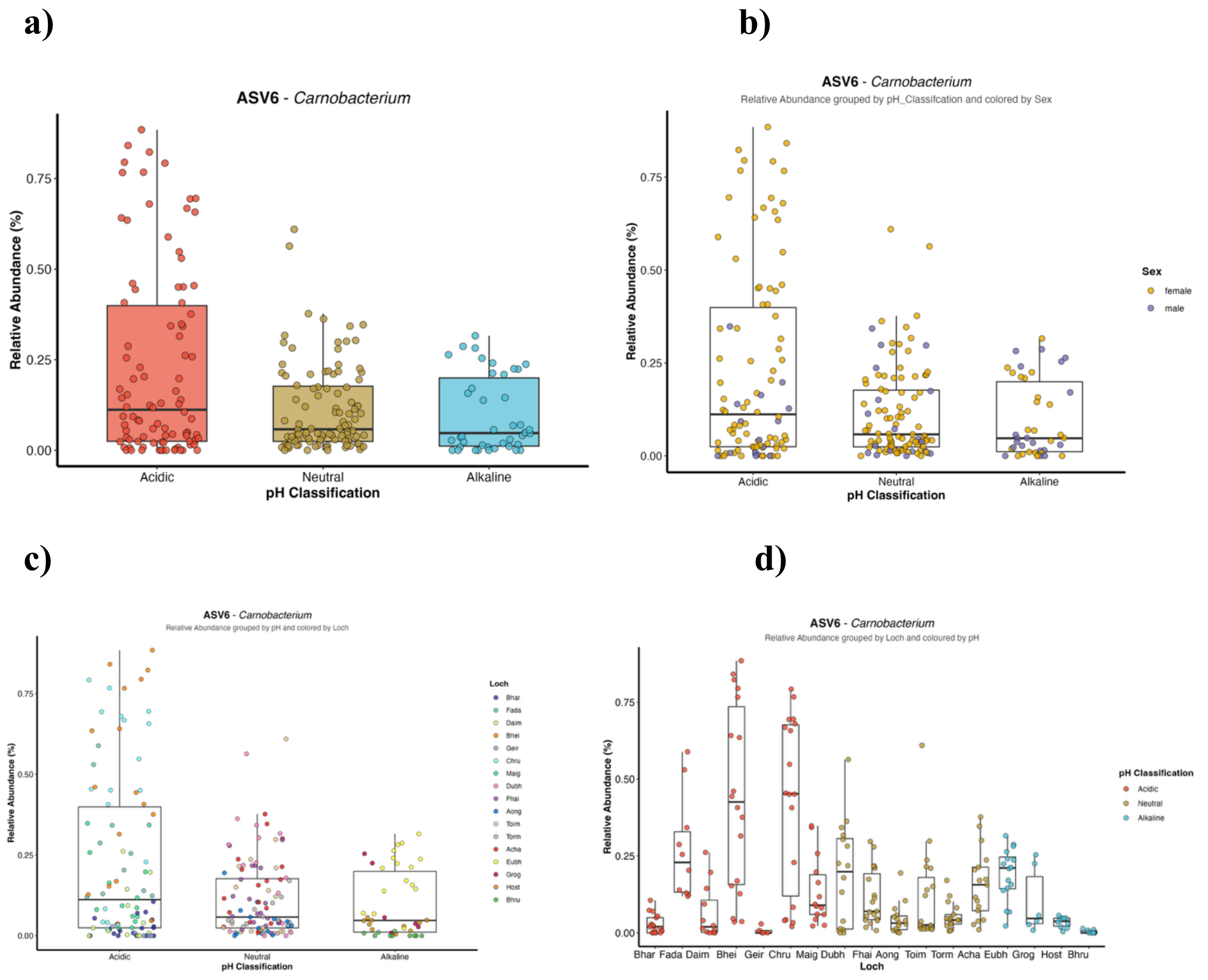


**Supplementary Figure 4:** Relative abundance of ASV6 (*Carnobacterium*) across pH classifications and host metadata. Relative abundance (%) of ASV6 grouped by: **(a)** pH classification (acidic, neutral, and alkaline); **(b)** pH classifications coloured by host sex (female and male); **(c)** pH classifications coloured by sampling location (loch). **(d)** Relative abundance of ASV6 across sampling locations (loch), with points coloured by pH classification (acidic, neutral, alkaline). Boxplots represent the median and interquartile range, with individual points showing sample-level variation.

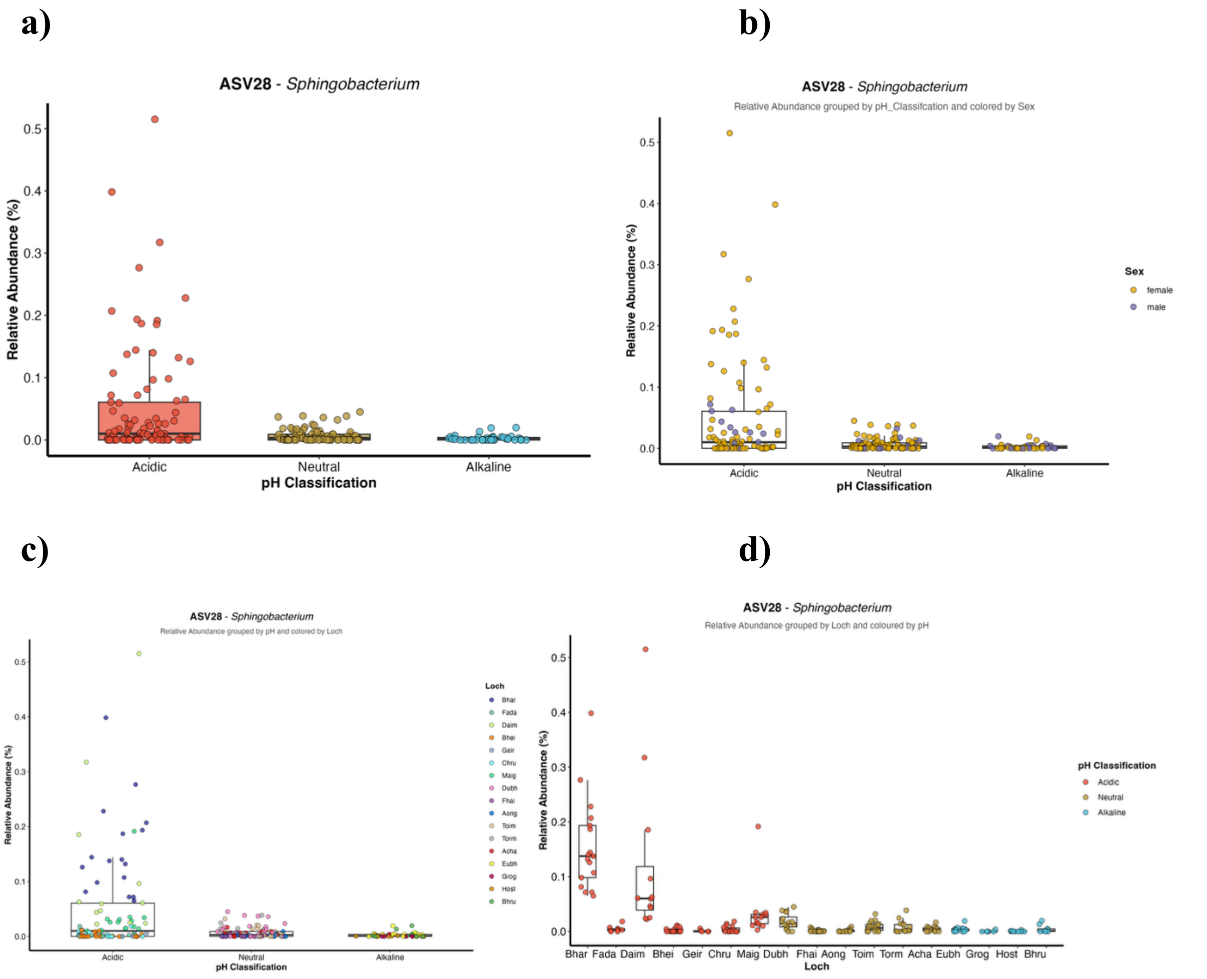


**Supplementary Figure 5:** Relative abundance of ASV28 (*Sphingobacterium*) across pH classifications and host metadata. Relative abundance (%) of ASV28 grouped by: **(a)** pH classification (acidic, neutral, and alkaline); **(b)** pH classifications coloured by host sex (female and male); **(c)** pH classifications coloured by sampling location (loch). **(d)** Relative abundance of ASV28 across sampling locations (loch), with points coloured by pH classification (acidic, neutral, alkaline). Boxplots represent the median and interquartile range, with individual points showing sample-level variation.

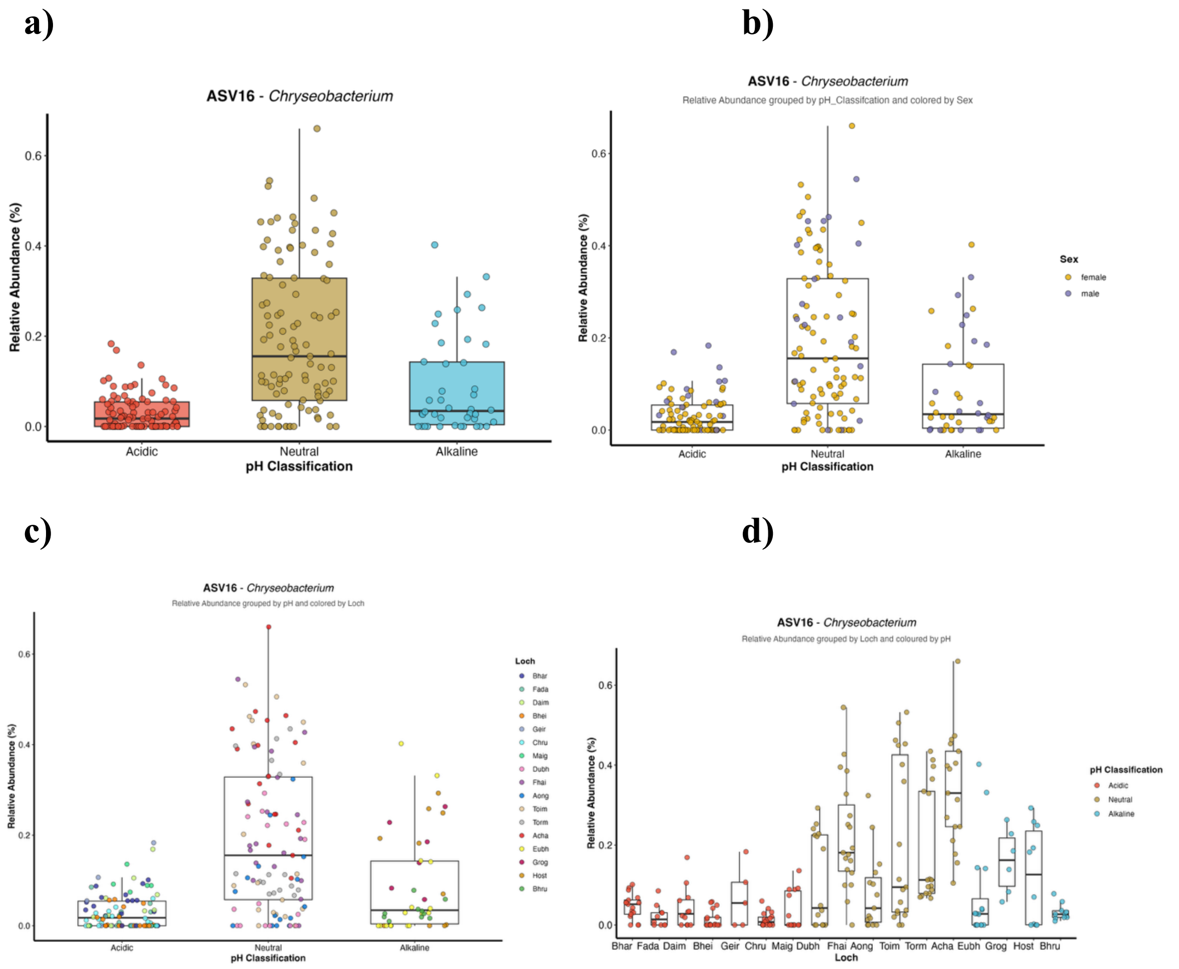


**Supplementary Figure 6:** Relative abundance of ASV16 (*Chryseobacterium*) across pH classifications and host metadata. Relative abundance (%) of ASV16 grouped by: **(a)** pH classification (acidic, neutral, and alkaline); **(b)** pH classifications coloured by host sex (female and male); **(c)** pH classifications coloured by sampling location (loch). **(d)** Relative abundance of ASV16 across sampling locations (loch), with points coloured by pH classification (acidic, neutral, alkaline). Boxplots represent the median and interquartile range, with individual points showing sample-level variation.

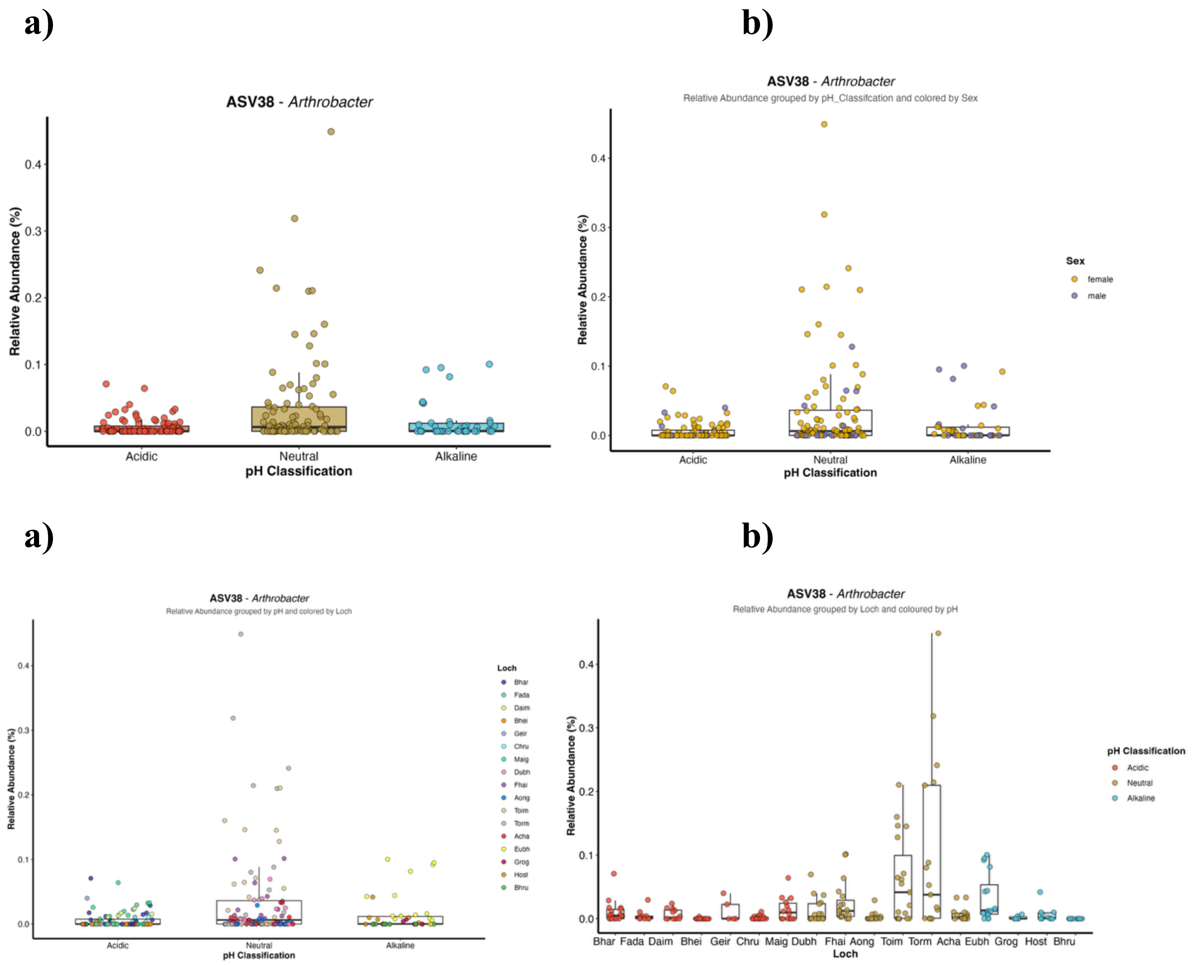


**Supplementary Figure 7:** Relative abundance of ASV38 (*Arthrobacter*) across pH classifications and host metadata. Relative abundance (%) of ASV28 grouped by: **(a)** pH classification (acidic, neutral, and alkaline); **(b)** pH classifications coloured by host sex (female and male); **(c)** pH classifications coloured by sampling location (loch). **(d)** Relative abundance of ASV38 across sampling locations (loch), with points coloured by pH classification (acidic, neutral, alkaline). Boxplots represent the median and interquartile range, with individual points showing sample-level variation.

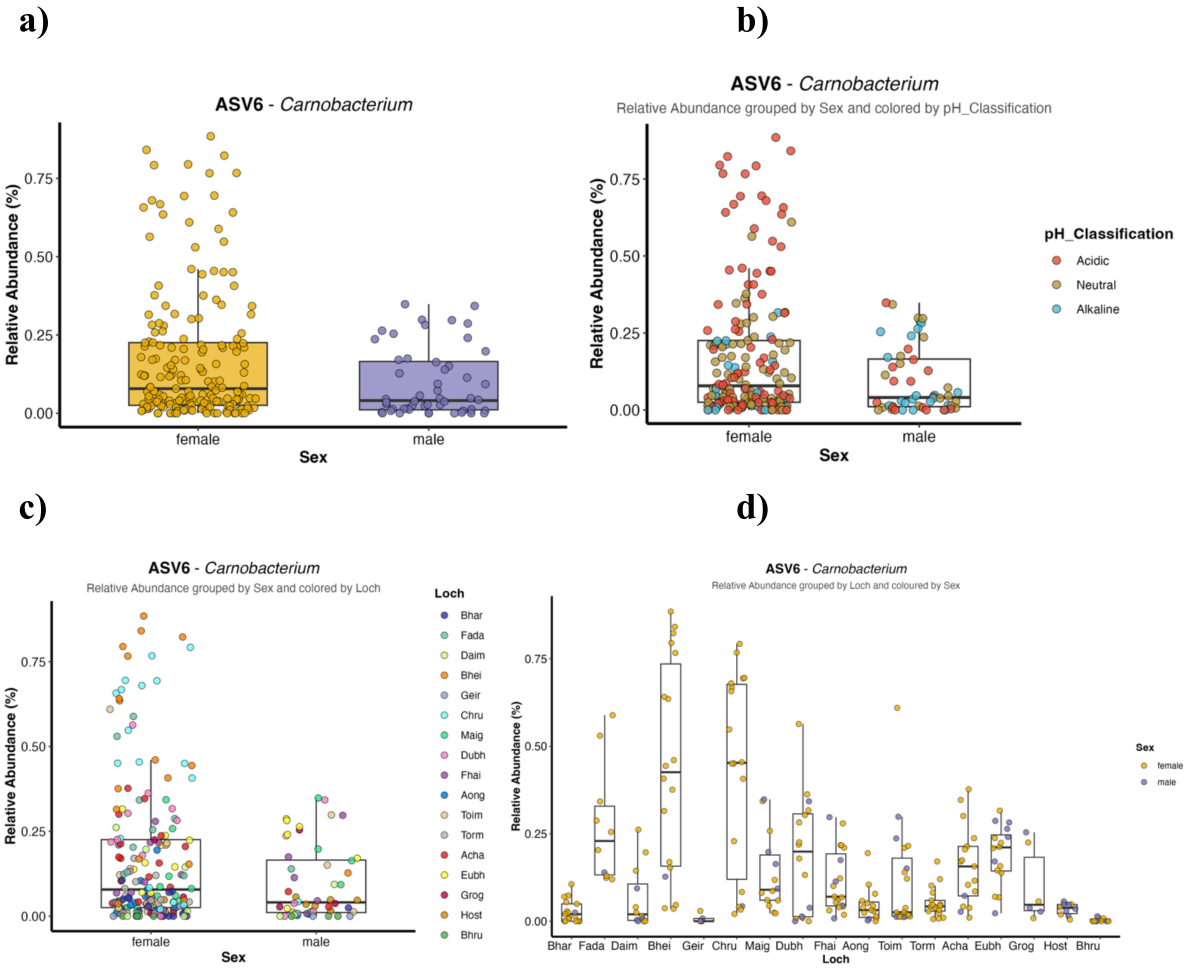


**Supplementary Figure 8:** Relative abundance of ASV6 (*Carnobacterium*) across host sex and environmental variables. Relative abundance (%) of ASV6 grouped by : **(a)** host sex (female and male); **(b)** host sex with points coloured by environmental pH classification (acidic, neutral, alkaline); **(c)** host sex with points coloured by sampling location (loch). **(d)** Relative abundance of ASV6 across sampling locations (loch), with points coloured by host sex (female and male). Boxplots represent the median and interquartile range, with individual points representing individual samples.

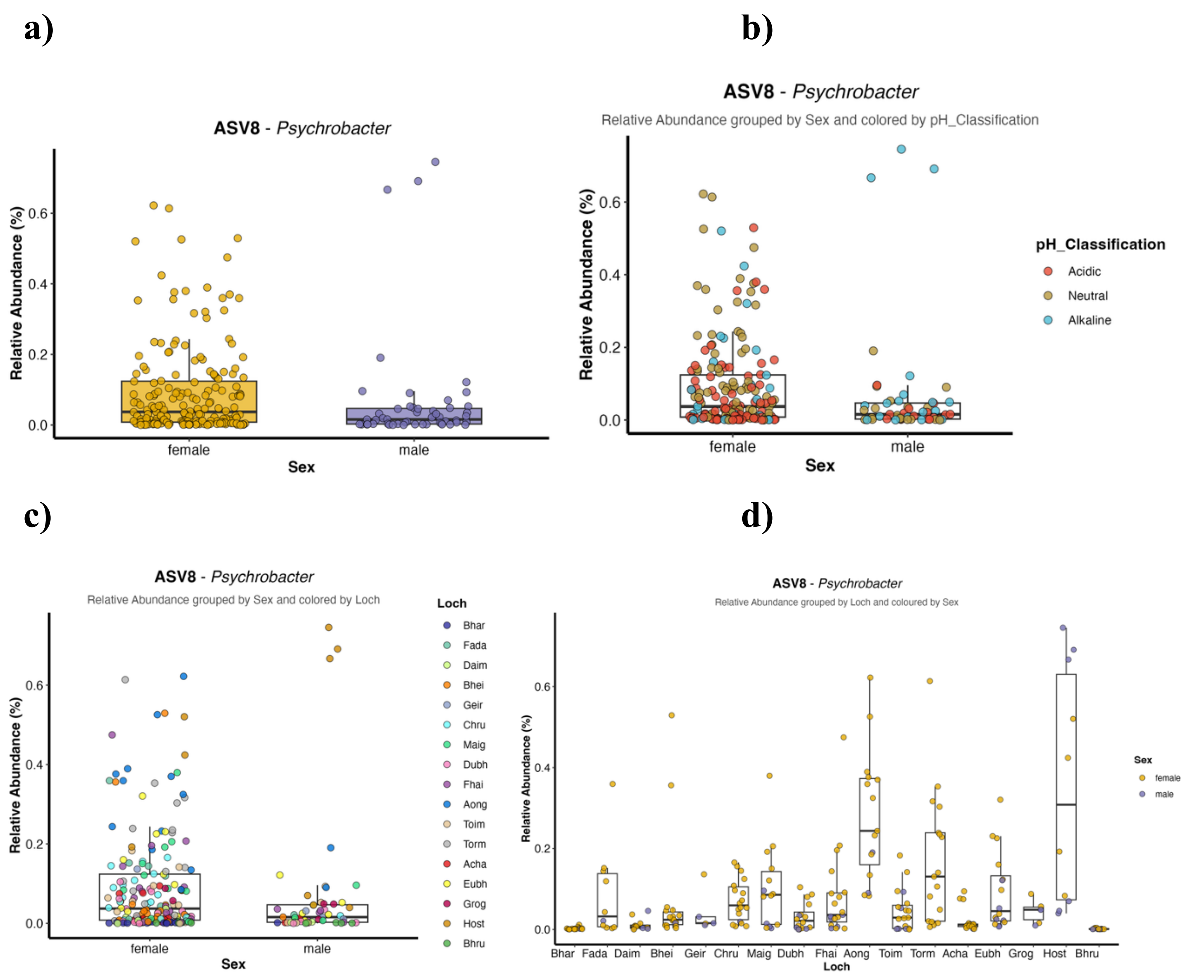


**Supplementary Figure 9:** Relative abundance of ASV8 (*Psychrobacter*) across host sex and environmental variables. Relative abundance (%) of ASV8 grouped by : **(a)** host sex (female and male); **(b)** host sex with points coloured by environmental pH classification (acidic, neutral, alkaline); **(c)** host sex with points coloured by sampling location (loch). **(d)** Relative abundance of ASV8 across sampling locations (loch), with points coloured by host sex (female and male). Boxplots represent the median and interquartile range, with individual points representing individual samples.

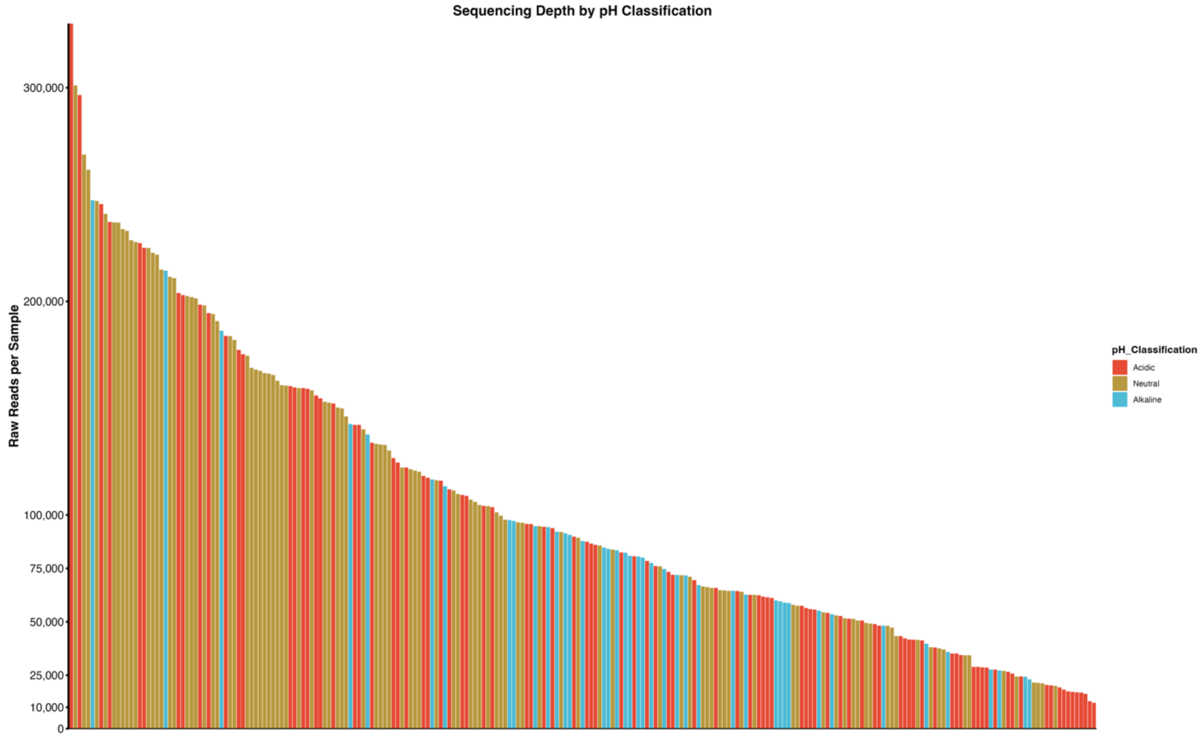


**Supplementary Figure 10:** Sequencing depth across samples grouped by environmental pH classification. Raw sequencing reads per sample are shown for all samples included in the dataset. Each bar represents an individual sample, ordered from highest to lowest sequencing depth. Bars are coloured according to environmental pH classification (acidic, neutral, and alkaline). This figure illustrates the distribution and variation in sequencing depth across samples and confirms that sequencing depths were comparable across pH categories.

| **Supplementary Table 1: Pairwise comparison among metadata variables.** | | |
|---|---|---|
| **Variable 1** | Variable 2 | Statistical test, p-value |
| **Sampling date** | Loch | Chiseq, p =2.2e-16 |
| **Sex** | Loch | Chiseq, p = 0.002 |
| **Sex** | Sampling date | Chiseq, p = 003 |
| **Length** | Loch | Aov, p = 0.6 |
| **Length** | Sex | Aov, p = 0.161 |
| **Copper (Cu)** | Zinc (Zn) | Cor, p = 0.104 |
| **Copper (Cu)** | Lead (Pb) | Cor, p = 0.578 |
| **pH** | Copper (CU) | Cor, p = 0.194 |
| **pH** | Conductivity | Cor, p = 9.955e-08 |

| **Supplementary Table 2: Summary of unique bacteria genera per loch** | | | | |
|---|---|---|---|---|
| **Loch** | Mean ± SD genera | Median genera | Lowest genera | Highest genera |
| Bhar | 19 ± 4 | 20 | 11 | 25 |
| Fada | 24 ± 8 | 22 | 14 | 43 |
| Daim | 17 ± 5 | 18 | 9 | 23 |
| Bhei | 16 ± 4 | 16 | 10 | 22 |
| Geir | 17 ± 5 | 17 | 11 | 23 |
| Chru | 19 ± 7 | 19 | 9 | 38 |
| Maig | 18 ± 4 | 17 | 13 | 23 |
| Dubh | 19 ± 13 | 17 | 8 | 64 |
| Fhai | 24 ± 7 | 23 | 13 | 41 |
| Aong | 24 ± 7 | 23 | 11 | 35 |
| Toim | 32 ± 15 | 25 | 16 | 61 |
| Torm | 30 ± 18 | 23 | 17 | 72 |
| Acha | 25 ± 4 | 24 | 19 | 37 |
| Eubh | 26 ± 7 | 24 | 17 | 40 |
| Grog | 22 ± 2 | 23 | 20 | 25 |
| Host | 26 ± 7 | 23 | 19 | 37 |
| Bhru | 17 ± 2 | 17 | 14 | 21 |

| Supplementary Table 3: Summary of bacterial genera abundance | | | | | |
|---|---|---|---|---|---|
| **Genus** | Mean Abundance (%) | Median Abundance (%) | Minimum Abundance (%) | Maximum Abundance (%) | samples present (%) |
| *Janthinobacterium* | 49.8699 | 52.3445 | 3.5119 | 91.0261 | 239 |
| *Pseudomonas* | 22.6959 | 20.1067 | 4.1928 | 62.7276 | 239 |
| *Acinetobacter* | 7.3553 | 2.9978 | 0.0750 | 53.3347 | 239 |
| *Psychrobacter* | 6.7415 | 2.1333 | 0.0098 | 76.1737 | 239 |
| *Exiguobacterium* | 4.4592 | 0.6241 | 0.0023 | 54.6014 | 219 |
| *Hydrogenophaga* | 2.6041 | 2.6041 | 0.0615 | 5.1466 | 2 |
| *Chryseobacterium* | 2.3795 | 1.3255 | 0.0267 | 16.6112 | 239 |
| *Carnobacterium* | 2.2110 | 0.8101 | 0.0062 | 31.7652 | 231 |
| *Comamonas* | 1.4368 | 1.0371 | 0.0397 | 5.8432 | 239 |
| *Cytobacillus* | 1.1558 | 0.0969 | 0.0014 | 4.4281 | 4 |
| *Arthrobacter* | 1.1091 | 0.1189 | 0.0015 | 21.9177 | 157 |
| *Rahnella* | 0.9281 | 0.1821 | 0.0077 | 10.0866 | 137 |
| *Brevundimonas* | 0.9175 | 0.0210 | 0.0025 | 35.7733 | 110 |
| *Aeromonas* | 0.3536 | 0.0595 | 0.0011 | 4.1782 | 117 |
| *Hydromonas* | 0.3358 | 0.0721 | 0.0005 | 2.4733 | 16 |
| *Flavobacterium* | 0.2917 | 0.0673 | 0.0036 | 12.1509 | 192 |
| *Cyanobium PCC-6307* | 0.2830 | 0.0369 | 0.0049 | 4.1911 | 65 |
| *Shewanella* | 0.2756 | 0.0982 | 0.0005 | 3.1428 | 156 |
| *Cypionkella* | 0.2414 | 0.2414 | 0.0623 | 0.4205 | 2 |
| *Mesobacillus* | 0.2040 | 0.0947 | 0.0145 | 1.3802 | 21 |
| *Staphylococcus* | 0.2031 | 0.0404 | 0.0004 | 6.1267 | 151 |
| *Stenotrophomonas* | 0.2028 | 0.0151 | 0.0015 | 7.8261 | 147 |
| *Deinococcus* | 0.1866 | 0.1866 | 0.1724 | 0.2009 | 2 |
| *Sphingobacterium* | 0.1791 | 0.0580 | 0.0026 | 2.6097 | 180 |
| *Enterococcus* | 0.1581 | 0.1142 | 0.0181 | 0.5516 | 10 |
| *Acidovorax* | 0.1572 | 0.1102 | 0.0047 | 1.4440 | 232 |
| *Deefgea* | 0.1444 | 0.0075 | 0.0004 | 1.8681 | 14 |
| *Lachnospiraceae NK4A136 group* | 0.1441 | 0.0103 | 0.0015 | 0.4204 | 3 |
| *Serratia* | 0.1311 | 0.0761 | 0.0089 | 2.8681 | 94 |
| *Bacteroides* | 0.1276 | 0.0062 | 0.0030 | 0.3738 | 3 |
| *Bacillus* | 0.1189 | 0.0168 | 0.0024 | 1.1049 | 12 |
| *Clostridium* | 0.1152 | 0.0178 | 0.0025 | 1.7307 | 32 |
| *Glutamicibacter* | 0.1087 | 0.1087 | 0.1087 | 0.1087 | 1 |
| *Prevotellaceae UCG-001* | 0.1065 | 0.1065 | 0.1065 | 0.1065 | 1 |
| *Curtobacterium* | 0.0945 | 0.0543 | 0.0080 | 0.3633 | 17 |
| *Turicimonas* | 0.0927 | 0.0927 | 0.0927 | 0.0927 | 1 |
| *Rhodoferax* | 0.0916 | 0.0143 | 0.0065 | 0.8857 | 12 |
| *Emticicia* | 0.0894 | 0.0894 | 0.0894 | 0.0894 | 1 |
| *Aurantimicrobium* | 0.0892 | 0.0404 | 0.0004 | 0.3364 | 27 |
| *Verticiella* | 0.0884 | 0.0104 | 0.0018 | 1.9313 | 49 |
| *Dubosiella* | 0.0818 | 0.0818 | 0.0818 | 0.0818 | 1 |
| *Pedobacter* | 0.0747 | 0.0250 | 0.0004 | 2.0876 | 167 |
| *Colidextribacter* | 0.0671 | 0.0671 | 0.0671 | 0.0671 | 1 |
| *alphaI cluster* | 0.0667 | 0.0163 | 0.0022 | 0.2319 | 4 |
| *Oscillibacter* | 0.0666 | 0.0666 | 0.0666 | 0.0666 | 1 |

| | | | | | |
|---|---|---|---|---|---|
| *Massilia* | 0.0629 | 0.0407 | 0.0028 | 0.3301 | 103 |
| *Citrobacter* | 0.0624 | 0.0338 | 0.0031 | 0.3691 | 70 |
| *Candidatus Gortzia* | 0.0613 | 0.0204 | 0.0007 | 0.1942 | 9 |
| *Intestinimonas* | 0.0599 | 0.0599 | 0.0599 | 0.0599 | 1 |
| *Butyribacter* | 0.0590 | 0.0590 | 0.0590 | 0.0590 | 1 |
| *CL500-29 marine group* | 0.0553 | 0.0162 | 0.0037 | 0.3040 | 15 |
| *Faecalibaculum* | 0.0531 | 0.0531 | 0.0196 | 0.0866 | 2 |
| *Planococcus* | 0.0495 | 0.0655 | 0.0013 | 0.0870 | 5 |
| *Alistipes* | 0.0485 | 0.0485 | 0.0485 | 0.0485 | 1 |
| *Muribaculum* | 0.0461 | 0.0461 | 0.0461 | 0.0461 | 1 |
| *Conexibacter* | 0.0454 | 0.0101 | 0.0019 | 0.3397 | 24 |
| *Ligilactobacillus* | 0.0442 | 0.0442 | 0.0442 | 0.0442 | 1 |
| *Pseudorhodobacter* | 0.0440 | 0.0440 | 0.0440 | 0.0440 | 1 |
| *Rhizobium* | 0.0438 | 0.0224 | 0.0077 | 0.1230 | 4 |
| *Mucispirillum* | 0.0428 | 0.0428 | 0.0428 | 0.0428 | 1 |
| *Candidatus Bacilloplasma* | 0.0424 | 0.0424 | 0.0262 | 0.0585 | 2 |
| *Desemzia* | 0.0402 | 0.0402 | 0.0402 | 0.0402 | 1 |
| *Parabacteroides* | 0.0380 | 0.0380 | 0.0380 | 0.0380 | 1 |
| *Lactococcus* | 0.0368 | 0.0083 | 0.0030 | 0.1812 | 8 |
| *Lachnoclostridium* | 0.0366 | 0.0366 | 0.0366 | 0.0366 | 1 |
| *hgcI clade* | 0.0357 | 0.0381 | 0.0096 | 0.0811 | 8 |
| *Acidiphilium* | 0.0352 | 0.0269 | 0.0021 | 0.1556 | 18 |
| *Fructilactobacillus* | 0.0350 | 0.0350 | 0.0350 | 0.0350 | 1 |
| *Legionella* | 0.0347 | 0.0302 | 0.0008 | 0.1256 | 28 |
| *Klenkia* | 0.0342 | 0.0165 | 0.0152 | 0.0708 | 3 |
| *Viridibacillus* | 0.0337 | 0.0337 | 0.0337 | 0.0337 | 1 |
| *Romboutsia* | 0.0330 | 0.0347 | 0.0047 | 0.0694 | 12 |
| *Salinicoccus* | 0.0322 | 0.0322 | 0.0322 | 0.0322 | 1 |
| *Zhengella* | 0.0321 | 0.0321 | 0.0321 | 0.0321 | 1 |
| *Pararhizobium* | 0.0306 | 0.0131 | 0.0018 | 0.1691 | 17 |
| *Sphingomonas* | 0.0303 | 0.0192 | 0.0019 | 0.1391 | 56 |
| *Rouxiella* | 0.0295 | 0.0295 | 0.0295 | 0.0295 | 1 |
| *Saccharedens* | 0.0282 | 0.0149 | 0.0030 | 0.1678 | 34 |
| *[Eubacterium] xylanophilum group* | 0.0281 | 0.0281 | 0.0015 | 0.0547 | 2 |
| *Lactobacillus* | 0.0281 | 0.0281 | 0.0124 | 0.0438 | 2 |
| *Undibacterium* | 0.0279 | 0.0047 | 0.0012 | 0.1011 | 4 |
| *Methylibium* | 0.0278 | 0.0278 | 0.0278 | 0.0278 | 1 |
| *Novosphingobium* | 0.0271 | 0.0233 | 0.0017 | 0.0835 | 26 |
| *Methylobacterium* | 0.0269 | 0.0192 | 0.0052 | 0.0581 | 6 |
| *Granulicatella* | 0.0268 | 0.0268 | 0.0027 | 0.0508 | 2 |
| *Mesorhizobium* | 0.0258 | 0.0258 | 0.0258 | 0.0258 | 1 |
| *Latilactobacillus* | 0.0257 | 0.0257 | 0.0257 | 0.0257 | 1 |
| *LD29* | 0.0253 | 0.0199 | 0.0154 | 0.0404 | 3 |
| *Lichenibacterium* | 0.0252 | 0.0139 | 0.0021 | 0.0972 | 12 |
| *Aerococcus* | 0.0242 | 0.0057 | 0.0034 | 0.1016 | 11 |
| *Cohnella* | 0.0242 | 0.0242 | 0.0242 | 0.0242 | 1 |
| *Blastocatella* | 0.0233 | 0.0233 | 0.0056 | 0.0410 | 2 |
| *Pantoea* | 0.0233 | 0.0233 | 0.0233 | 0.0233 | 1 |
| *Hyphomicrobium* | 0.0229 | 0.0155 | 0.0021 | 0.0705 | 11 |

| | | | | | |
|---|---|---|---|---|---|
| *Peribacillus* | 0.0223 | 0.0223 | 0.0223 | 0.0223 | 1 |
| *Variovorax* | 0.0221 | 0.0192 | 0.0026 | 0.0624 | 40 |
| *Caedibacter* | 0.0219 | 0.0124 | 0.0087 | 0.0537 | 6 |
| *Enhydrobacter* | 0.0216 | 0.0058 | 0.0033 | 0.0762 | 11 |
| *Niallia* | 0.0215 | 0.0121 | 0.0038 | 0.0484 | 3 |
| *Acutalibacter* | 0.0214 | 0.0214 | 0.0214 | 0.0214 | 1 |
| *Finegoldia* | 0.0212 | 0.0212 | 0.0212 | 0.0212 | 1 |
| *Candidatus Paracaedibacter* | 0.0212 | 0.0212 | 0.0212 | 0.0212 | 1 |
| *Agathobaculum* | 0.0209 | 0.0209 | 0.0209 | 0.0209 | 1 |
| *Empedobacter* | 0.0207 | 0.0207 | 0.0207 | 0.0207 | 1 |
| *Limnohabitans* | 0.0203 | 0.0073 | 0.0026 | 0.0809 | 7 |
| *Polynucleobacter* | 0.0201 | 0.0106 | 0.0008 | 0.1654 | 85 |
| *Polaromonas* | 0.0193 | 0.0062 | 0.0025 | 0.0624 | 4 |
| *Turicibacter* | 0.0192 | 0.0063 | 0.0058 | 0.0628 | 5 |
| *Candidatus Megaira* | 0.0190 | 0.0076 | 0.0021 | 0.0604 | 5 |
| *Aureimonas* | 0.0186 | 0.0177 | 0.0040 | 0.0351 | 4 |
| *Odoribacter* | 0.0185 | 0.0185 | 0.0185 | 0.0185 | 1 |
| *Luteimicrobium* | 0.0184 | 0.0184 | 0.0184 | 0.0184 | 1 |
| *Candidatus Branchiomonas* | 0.0183 | 0.0076 | 0.0018 | 0.0807 | 9 |
| *Fenollaria* | 0.0182 | 0.0182 | 0.0182 | 0.0182 | 1 |
| *Mycobacterium* | 0.0175 | 0.0063 | 0.0005 | 0.0852 | 37 |
| *Frondihabitans* | 0.0175 | 0.0150 | 0.0028 | 0.0371 | 4 |
| *FukuN18 freshwater group* | 0.0172 | 0.0124 | 0.0022 | 0.0429 | 12 |
| *Dyadobacter* | 0.0172 | 0.0172 | 0.0158 | 0.0187 | 2 |
| *Trichococcus* | 0.0172 | 0.0199 | 0.0067 | 0.0225 | 4 |
| *Agrococcus* | 0.0172 | 0.0096 | 0.0006 | 0.0632 | 10 |
| *Methylorosula* | 0.0170 | 0.0208 | 0.0025 | 0.0293 | 5 |
| *Tardiphaga* | 0.0169 | 0.0077 | 0.0056 | 0.0466 | 4 |
| *Candidatus Trichorickettsia* | 0.0166 | 0.0147 | 0.0034 | 0.0384 | 7 |
| *Buttiauxella* | 0.0166 | 0.0166 | 0.0131 | 0.0201 | 2 |
| *Gloeobacter PCC-7421* | 0.0161 | 0.0161 | 0.0103 | 0.0218 | 2 |
| *Duganella* | 0.0160 | 0.0087 | 0.0008 | 0.0661 | 34 |
| *Paenarthrobacter* | 0.0160 | 0.0160 | 0.0012 | 0.0308 | 2 |
| *Cellulosilyticum* | 0.0160 | 0.0160 | 0.0059 | 0.0260 | 2 |
| *Lachnospiraceae FCS020 group* | 0.0160 | 0.0160 | 0.0010 | 0.0309 | 2 |
| *Streptococcus* | 0.0158 | 0.0126 | 0.0023 | 0.0449 | 11 |
| *Aliivibrio* | 0.0155 | 0.0155 | 0.0155 | 0.0155 | 1 |
| *Iodobacter* | 0.0153 | 0.0110 | 0.0004 | 0.0804 | 42 |
| *Hymenobacter* | 0.0153 | 0.0183 | 0.0012 | 0.0344 | 12 |
| *Chryseomicrobium* | 0.0150 | 0.0150 | 0.0150 | 0.0150 | 1 |
| *Nocardioides* | 0.0146 | 0.0146 | 0.0025 | 0.0266 | 2 |
| *Paeniclostridium* | 0.0145 | 0.0145 | 0.0050 | 0.0240 | 2 |
| *Paracoccus* | 0.0142 | 0.0139 | 0.0026 | 0.0330 | 10 |
| *HT002* | 0.0141 | 0.0141 | 0.0054 | 0.0228 | 2 |
| *Tabrizicola* | 0.0140 | 0.0168 | 0.0070 | 0.0183 | 3 |
| *Gemmata* | 0.0140 | 0.0140 | 0.0100 | 0.0180 | 2 |
| *Rugamonas* | 0.0138 | 0.0077 | 0.0017 | 0.0463 | 20 |
| *Rickettsiella* | 0.0134 | 0.0115 | 0.0014 | 0.0557 | 14 |
| *Roseiarcus* | 0.0131 | 0.0060 | 0.0006 | 0.0436 | 11 |
| *Rubritepida* | 0.0130 | 0.0130 | 0.0130 | 0.0130 | 1 |

| | | | | | |
|---|---|---|---|---|---|
| *Pajaroellobacter* | 0.0129 | 0.0129 | 0.0129 | 0.0129 | 1 |
| *Leucobacter* | 0.0127 | 0.0056 | 0.0027 | 0.0452 | 6 |
| *Marisediminicola* | 0.0127 | 0.0127 | 0.0127 | 0.0127 | 1 |
| *Paeniglutamicibacter* | 0.0126 | 0.0103 | 0.0059 | 0.0312 | 7 |
| *Enteractinococcus* | 0.0126 | 0.0126 | 0.0126 | 0.0126 | 1 |
| *Microterricola* | 0.0124 | 0.0124 | 0.0124 | 0.0124 | 1 |
| *Kocuria* | 0.0123 | 0.0115 | 0.0073 | 0.0189 | 4 |
| *Methylocystis* | 0.0121 | 0.0117 | 0.0033 | 0.0218 | 4 |
| *Rhodopseudomonas* | 0.0121 | 0.0121 | 0.0121 | 0.0121 | 1 |
| *FukuN57* | 0.0120 | 0.0120 | 0.0120 | 0.0120 | 1 |
| *Eisenbergiella* | 0.0119 | 0.0119 | 0.0119 | 0.0119 | 1 |
| *Ruminiclostridium* | 0.0112 | 0.0112 | 0.0076 | 0.0147 | 2 |
| *Lysobacter* | 0.0111 | 0.0111 | 0.0111 | 0.0111 | 1 |
| *Aquicella* | 0.0110 | 0.0088 | 0.0020 | 0.0304 | 10 |
| *Alkanindiges* | 0.0109 | 0.0109 | 0.0109 | 0.0109 | 1 |
| *Agrobacterium* | 0.0107 | 0.0081 | 0.0013 | 0.0337 | 12 |
| *Methylocella* | 0.0105 | 0.0060 | 0.0025 | 0.0230 | 3 |
| *Anaerotruncus* | 0.0105 | 0.0105 | 0.0105 | 0.0105 | 1 |
| *Candidatus Planktoluna* | 0.0103 | 0.0063 | 0.0031 | 0.0262 | 7 |
| *Allorhizobium* | 0.0102 | 0.0102 | 0.0102 | 0.0102 | 1 |
| *Eperythrozoon* | 0.0102 | 0.0102 | 0.0045 | 0.0159 | 2 |
| *Candidatus Bealeia* | 0.0102 | 0.0103 | 0.0053 | 0.0147 | 4 |
| *Alsobacter* | 0.0101 | 0.0101 | 0.0101 | 0.0101 | 1 |
| *Family XIII AD3011 group* | 0.0101 | 0.0101 | 0.0101 | 0.0101 | 1 |
| *GCA-900066575* | 0.0100 | 0.0100 | 0.0100 | 0.0100 | 1 |
| *Mobilitalea* | 0.0099 | 0.0099 | 0.0099 | 0.0099 | 1 |
| *Candidatus Udaeobacter* | 0.0099 | 0.0099 | 0.0099 | 0.0099 | 1 |
| *Phormidium SAG 37.90* | 0.0099 | 0.0099 | 0.0099 | 0.0099 | 1 |
| *Reyranella* | 0.0096 | 0.0096 | 0.0096 | 0.0096 | 1 |
| *UBA12409* | 0.0095 | 0.0068 | 0.0028 | 0.0238 | 5 |
| *Paenibacillus* | 0.0094 | 0.0037 | 0.0005 | 0.0504 | 26 |
| *Mucilaginibacter* | 0.0094 | 0.0094 | 0.0094 | 0.0094 | 1 |
| *Achromobacter* | 0.0093 | 0.0093 | 0.0093 | 0.0093 | 1 |
| *Blastococcus* | 0.0091 | 0.0091 | 0.0036 | 0.0145 | 2 |
| *Burkholderia-Caballeronia-Paraburkholderia* | 0.0089 | 0.0092 | 0.0009 | 0.0248 | 12 |
| *Corynebacterium* | 0.0089 | 0.0036 | 0.0004 | 0.0696 | 13 |
| *Terriglobus* | 0.0088 | 0.0088 | 0.0088 | 0.0088 | 1 |
| *Fonticella* | 0.0088 | 0.0088 | 0.0088 | 0.0088 | 1 |
| *Tyzzerella* | 0.0088 | 0.0088 | 0.0088 | 0.0088 | 1 |
| *Peptoniphilus* | 0.0088 | 0.0088 | 0.0088 | 0.0088 | 1 |
| *Yersinia* | 0.0084 | 0.0081 | 0.0008 | 0.0201 | 8 |
| *Paludibaculum* | 0.0083 | 0.0083 | 0.0083 | 0.0083 | 1 |
| *Labrys* | 0.0081 | 0.0055 | 0.0023 | 0.0193 | 4 |
| *Fuscovulum* | 0.0081 | 0.0081 | 0.0061 | 0.0100 | 2 |
| *Delftia* | 0.0079 | 0.0053 | 0.0009 | 0.0217 | 7 |
| *Luteimonas* | 0.0079 | 0.0079 | 0.0079 | 0.0079 | 1 |
| *Helcococcus* | 0.0077 | 0.0077 | 0.0077 | 0.0077 | 1 |
| *Bryobacter* | 0.0075 | 0.0075 | 0.0075 | 0.0075 | 1 |
| *Roseomonas* | 0.0074 | 0.0074 | 0.0028 | 0.0119 | 2 |

| | | | | | |
|---|---|---|---|---|---|
| *Roseburia* | 0.0073 | 0.0073 | 0.0073 | 0.0073 | 1 |
| *Gemmiger* | 0.0073 | 0.0073 | 0.0073 | 0.0073 | 1 |
| *Abiotrophia* | 0.0073 | 0.0073 | 0.0073 | 0.0073 | 1 |
| *Rhizorhabdus* | 0.0072 | 0.0072 | 0.0072 | 0.0072 | 1 |
| *Epulopiscium* | 0.0072 | 0.0072 | 0.0072 | 0.0072 | 1 |
| *Huanghella* | 0.0072 | 0.0072 | 0.0072 | 0.0072 | 1 |
| *Pirellula* | 0.0071 | 0.0071 | 0.0071 | 0.0071 | 1 |
| *Rothia* | 0.0070 | 0.0093 | 0.0006 | 0.0111 | 3 |
| *Caulobacter* | 0.0069 | 0.0069 | 0.0046 | 0.0092 | 2 |
| *Pedomicrobium* | 0.0069 | 0.0069 | 0.0045 | 0.0093 | 2 |
| *Azorhizobium* | 0.0068 | 0.0068 | 0.0051 | 0.0086 | 2 |
| *Pasteuria* | 0.0068 | 0.0068 | 0.0068 | 0.0068 | 1 |
| *Psychrobacillus* | 0.0067 | 0.0019 | 0.0018 | 0.0164 | 3 |
| *Coxiella* | 0.0067 | 0.0055 | 0.0031 | 0.0126 | 4 |
| *ASF356* | 0.0067 | 0.0067 | 0.0067 | 0.0067 | 1 |
| *Marvinbryantia* | 0.0067 | 0.0067 | 0.0067 | 0.0067 | 1 |
| *Candidatus Solibacter* | 0.0063 | 0.0063 | 0.0063 | 0.0063 | 1 |
| *UCG-005* | 0.0062 | 0.0062 | 0.0062 | 0.0062 | 1 |
| *[Ruminococcus] gauvreauii group* | 0.0062 | 0.0062 | 0.0062 | 0.0062 | 1 |
| *Segatella* | 0.0062 | 0.0057 | 0.0025 | 0.0103 | 3 |
| *Pseudarthrobacter* | 0.0061 | 0.0061 | 0.0061 | 0.0061 | 1 |
| *Fluviicola* | 0.0060 | 0.0060 | 0.0060 | 0.0060 | 1 |
| *Terrimicrobium* | 0.0059 | 0.0070 | 0.0011 | 0.0096 | 3 |
| *Pseudoxanthomonas* | 0.0059 | 0.0036 | 0.0007 | 0.0226 | 18 |
| *Catellicoccus* | 0.0058 | 0.0058 | 0.0058 | 0.0058 | 1 |
| *Paucibacter* | 0.0058 | 0.0043 | 0.0007 | 0.0186 | 10 |
| *Nubsella* | 0.0056 | 0.0056 | 0.0056 | 0.0056 | 1 |
| *Polymorphobacter* | 0.0055 | 0.0055 | 0.0030 | 0.0080 | 2 |
| *Anaerovorax* | 0.0055 | 0.0055 | 0.0055 | 0.0055 | 1 |
| *Kytococcus* | 0.0054 | 0.0029 | 0.0020 | 0.0145 | 5 |
| *Aliterella* | 0.0053 | 0.0053 | 0.0051 | 0.0055 | 2 |
| *Veillonella* | 0.0052 | 0.0057 | 0.0009 | 0.0091 | 3 |
| *Azospirillum* | 0.0051 | 0.0051 | 0.0051 | 0.0051 | 1 |
| *Candidatus Nitrosoarchaeum* | 0.0050 | 0.0050 | 0.0050 | 0.0050 | 1 |
| *Fastidiosipila* | 0.0050 | 0.0050 | 0.0050 | 0.0050 | 1 |
| *Oribacterium* | 0.0048 | 0.0048 | 0.0048 | 0.0048 | 1 |
| *MWH-Ta3* | 0.0048 | 0.0048 | 0.0042 | 0.0054 | 2 |
| *Baekduia* | 0.0048 | 0.0048 | 0.0048 | 0.0048 | 1 |
| *Devosia* | 0.0047 | 0.0047 | 0.0029 | 0.0066 | 2 |
| *Nitrospira* | 0.0047 | 0.0047 | 0.0022 | 0.0072 | 2 |
| *RS62 marine group* | 0.0046 | 0.0046 | 0.0040 | 0.0051 | 2 |
| *Noviherbaspirillum* | 0.0045 | 0.0041 | 0.0016 | 0.0079 | 6 |
| *Bryocella* | 0.0045 | 0.0045 | 0.0045 | 0.0045 | 1 |
| *Candidatus Endonucleariobacter* | 0.0045 | 0.0045 | 0.0045 | 0.0045 | 1 |
| *Saccharospirillum* | 0.0045 | 0.0045 | 0.0045 | 0.0045 | 1 |
| *JGI 0001001-H03* | 0.0044 | 0.0044 | 0.0044 | 0.0044 | 1 |
| *Paenochrobactrum* | 0.0044 | 0.0044 | 0.0044 | 0.0044 | 1 |
| *Thermomonas* | 0.0043 | 0.0043 | 0.0043 | 0.0043 | 1 |
| *Adlercreutzia* | 0.0043 | 0.0043 | 0.0043 | 0.0043 | 1 |

| | | | | | |
|---|---|---|---|---|---|
| *Thomasclavelia* | 0.0043 | 0.0043 | 0.0043 | 0.0043 | 1 |
| *Candidatus Ovatusbacter* | 0.0041 | 0.0041 | 0.0034 | 0.0048 | 2 |
| *Leuconostoc* | 0.0040 | 0.0040 | 0.0040 | 0.0040 | 1 |
| *Streptomyces* | 0.0036 | 0.0036 | 0.0036 | 0.0036 | 1 |
| *Alkalibacterium* | 0.0035 | 0.0035 | 0.0035 | 0.0035 | 1 |
| *Verruc-01* | 0.0034 | 0.0034 | 0.0034 | 0.0034 | 1 |
| *Acetatifactor* | 0.0033 | 0.0033 | 0.0033 | 0.0033 | 1 |
| *Enterocloster* | 0.0033 | 0.0033 | 0.0033 | 0.0033 | 1 |
| *Sporichthya* | 0.0033 | 0.0033 | 0.0033 | 0.0033 | 1 |
| *Occallatibacter* | 0.0032 | 0.0032 | 0.0032 | 0.0032 | 1 |
| *Amnibacterium* | 0.0031 | 0.0031 | 0.0031 | 0.0031 | 1 |
| *Frigoribacterium* | 0.0031 | 0.0031 | 0.0031 | 0.0031 | 1 |
| *Arcicella* | 0.0028 | 0.0028 | 0.0028 | 0.0028 | 1 |
| *Candidatus Xiphinematobacter* | 0.0028 | 0.0028 | 0.0015 | 0.0042 | 2 |
| *Escherichia-Shigella* | 0.0027 | 0.0027 | 0.0027 | 0.0027 | 1 |
| *Gaiella* | 0.0027 | 0.0027 | 0.0027 | 0.0027 | 1 |
| *Angustibacter* | 0.0025 | 0.0025 | 0.0025 | 0.0025 | 1 |
| *Jeotgalibacillus* | 0.0025 | 0.0025 | 0.0025 | 0.0025 | 1 |
| *Rhodococcus* | 0.0024 | 0.0030 | 0.0012 | 0.0033 | 5 |
| *Asticcacaulis* | 0.0024 | 0.0024 | 0.0015 | 0.0033 | 2 |
| *Bifidobacterium* | 0.0024 | 0.0024 | 0.0024 | 0.0024 | 1 |
| *Blautia* | 0.0024 | 0.0024 | 0.0024 | 0.0024 | 1 |
| *Arenimonas* | 0.0023 | 0.0023 | 0.0009 | 0.0038 | 2 |
| *Rhodovarius* | 0.0023 | 0.0023 | 0.0023 | 0.0023 | 1 |
| *Lautropia* | 0.0022 | 0.0022 | 0.0022 | 0.0022 | 1 |
| *Synechococcus PCC-7502* | 0.0021 | 0.0021 | 0.0021 | 0.0021 | 1 |
| *Brevibacterium* | 0.0021 | 0.0023 | 0.0012 | 0.0027 | 3 |
| *Parageobacillus* | 0.0020 | 0.0020 | 0.0020 | 0.0020 | 1 |
| *Bilophila* | 0.0019 | 0.0019 | 0.0019 | 0.0019 | 1 |
| *Pigmentiphaga* | 0.0018 | 0.0012 | 0.0005 | 0.0042 | 13 |
| *Breznakia* | 0.0017 | 0.0017 | 0.0017 | 0.0017 | 1 |
| *MM1* | 0.0016 | 0.0016 | 0.0014 | 0.0019 | 2 |
| *Aquihabitans* | 0.0015 | 0.0015 | 0.0015 | 0.0015 | 1 |
| *Sanguibacter* | 0.0015 | 0.0015 | 0.0015 | 0.0015 | 2 |
| *Phreatobacter* | 0.0014 | 0.0014 | 0.0014 | 0.0014 | 1 |
| *Skermanella* | 0.0012 | 0.0012 | 0.0012 | 0.0012 | 1 |
| *Actinomycetospora* | 0.0012 | 0.0012 | 0.0012 | 0.0012 | 1 |
| *Sporosarcina* | 0.0011 | 0.0011 | 0.0011 | 0.0011 | 1 |
| *Hirschia* | 0.0011 | 0.0011 | 0.0011 | 0.0011 | 1 |
| *Sphaerotilus* | 0.0009 | 0.0009 | 0.0009 | 0.0009 | 1 |
| *Cellulosimicrobium* | 0.0008 | 0.0008 | 0.0008 | 0.0008 | 1 |
| *Gordonia* | 0.0005 | 0.0005 | 0.0005 | 0.0005 | 1 |
| *Micrococcus* | 0.0004 | 0.0004 | 0.0004 | 0.0004 | 1 |

| Supplementary Table 4: Differentially abundant ASVs across pH classes identified using DESeq2 | | | | |
|---|---|---|---|---|
| **ASV_ID** | **log2FoldChange** | **p-value adjusted** | **Genus** | **Mean Abundance (%)** |
| ASV8 | 1.83 | 0.002678911 | Psychrobacter | 22.7194 |
| ASV22 | -0.84 | 0.011745347 | Pseudomonas | 17.3379 |
| ASV6 | -1.48 | 0.041169374 | Carnobacterium | 15.5822 |
| ASV13 | 2.07 | 0.009953592 | Exiguobacterium | 11.6088 |
| ASV16 | 2.47 | 5.20415E-05 | Chryseobacterium | 11.1735 |
| ASV37 | 1.26 | 0.048061675 | Chryseobacterium | 6.0139 |
| ASV65 | 1.46 | 0.046813345 | Acinetobacter | 2.6073 |
| ASV28 | -2.32 | 0.00104302 | Sphingobacterium | 2.3411 |
| ASV44 | 4.20 | 0.004114555 | Aeromonas | 2.2612 |
| ASV38 | 2.61 | 0.022741583 | Arthrobacter | 2.0695 |
| ASV88 | 1.85 | 0.045178987 | Acinetobacter | 1.3136 |
| ASV49 | 1.93 | 0.006607591 | Psychrobacter | 1.0677 |
| ASV42 | 4.19 | 3.70229E-08 | Acinetobacter | 0.9876 |
| ASV164 | 1.17 | 0.03488729 | Pseudomonas | 0.7831 |
| ASV120 | 3.99 | 0.002719379 | Acinetobacter | 0.4225 |
| ASV121 | 24.96 | 4.04488E-08 | Chryseobacterium | 0.3069 |
| ASV118 | 11.56 | 1.83112E-13 | Cyanobium PCC-6307 | 0.2379 |
| ASV97 | 24.58 | 6.55375E-21 | Arthrobacter | 0.2359 |
| ASV60 | 2.31 | 0.045178987 | Acinetobacter | 0.2113 |
| ASV167 | 4.01 | 0.045178987 | Chryseobacterium | 0.1346 |
| ASV124 | -16.29 | 7.18409E-15 | Cyanobacteria | 0.1315 |
| ASV163 | 27.22 | 1.47321E-34 | Cyanobium PCC-6307 | 0.1126 |
| ASV156 | 4.10 | 0.046813345 | Planctomycetes | 0.0979 |
| ASV242 | -24.11 | 6.88241E-29 | Cyanobacteria | 0.0864 |
| ASV333 | 2.05 | 0.045178987 | Acinetobacter | 0.0809 |
| ASV235 | 23.33 | 2.43539E-22 | Arthrobacter | 0.0388 |
| ASV248 | 24.67 | 5.496E-08 | Cyanobium PCC-6307 | 0.0360 |

Amplicon sequence variants (ASVs) showing significant differential abundance across pH classes were identified using DESeq2. The table includes ASV identifiers, log2 fold-change values, adjusted P-values (Benjamini–Hochberg correction), taxonomic genus assignments, and mean relative abundance across samples. Positive log2 fold change values indicate higher abundance in neutral/alkaline pH classes relative to acidic systems, whereas negative values indicate enrichment in acidic lochs. ASV124, ASV156 and AVS242 are phylum names as those ASVs were not assigned their genera name.

| Supplementary Table 5: Differentially abundant ASVs between host sex identified using DESeq2 | | | | |
|---|---|---|---|---|
| **ASV_ID** | **log2FoldChange** | **p-value adjusted** | **Genus** | **Mean Abundance (%)** |
| ASV3 | 0.198 | 0.018572948 | Janthinobacterium | 29.2474 |
| ASV4 | 0.232 | 0.013120249 | Janthinobacterium | 25.9847 |
| ASV8 | -1.526 | 0.006196673 | Psychrobacter | 8.7418 |
| ASV7 | -0.927 | 0.0127086 | Pseudomonas | 6.2564 |
| ASV5 | -1.008 | 0.014394508 | Pseudomonas | 6.1536 |
| ASV6 | -1.194 | 0.045357655 | Carnobacterium | 5.3005 |
| ASV23 | -0.468 | 0.012409471 | Janthinobacterium | 4.2725 |
| ASV19 | -1.721 | 0.030019951 | Exiguobacterium | 3.9000 |
| ASV27 | -0.577 | 0.006196673 | Janthinobacterium | 3.6721 |
| ASV11 | -1.456 | 0.004486665 | Psychrobacter | 2.7571 |
| ASV34 | -0.706 | 0.014394508 | Pseudomonas | 1.1649 |
| ASV31 | -0.884 | 0.025578056 | Pseudomonas | 1.1481 |
| ASV28 | -1.427 | 0.042669217 | Sphingobacterium | 0.3606 |
| ASV84 | -8.201 | 3.08213E-13 | Staphylococcus | 0.2774 |
| ASV35 | -0.512 | 0.049211764 | Pseudomonas | 0.2530 |
| ASV98 | -2.698 | 0.0263413 | Pseudomonas | 0.1907 |
| ASV51 | -0.775 | 0.0263413 | Pseudomonas | 0.1382 |
| ASV146 | -2.229 | 0.0205029 | Comamonas | 0.0682 |
| ASV171 | -5.121 | 0.014394508 | Verticiella | 0.0653 |
| ASV366 | -2.651 | 0.014394508 | Acidovorax | 0.0175 |
| ASV380 | -3.556 | 0.018572948 | Chryseobacterium | 0.0106 |
| ASV391 | -2.188 | 0.021069979 | Massilia | 0.0088 |
| ASV426 | -3.454 | 0.000258152 | Acinetobacter | 0.0070 |
| ASV545 | -2.350 | 0.014394508 | Acinetobacter | 0.0033 |

Amplicon sequence variants (ASVs) showing significant differential abundance between male and female hosts were identified using DESeq2. The table includes ASV identifiers, log2 fold-change values, adjusted P-values (Benjamini–Hochberg correction), genus-level taxonomic assignments, and mean relative abundance across samples. Positive log2 fold change values indicate enrichment in males, whereas negative values indicate enrichment in females.
